# A funnel approach to enable analyses of epitope-specific human CD4 T cells specific for influenza and SARS-CoV-2

**DOI:** 10.1101/2025.10.16.682991

**Authors:** K.A. Richards, R.C. Mettelman, K. Trombley, E.K. Allen, Z.B. Scott, A. Joachimiak, S. Schultz-Cherry, F. A. Chaves, P.G. Thomas, A.J. Sant

## Abstract

Protection from pathogenic organisms relies heavily on the adaptive immune response, for which key regulators are CD4 T cells. CD4 T cells, notable in the complexity of their repertoire and functional potential, can most easily be dissected with the ability to identify, quantify, characterize and isolate epitope-specific cells. In the study reported here, we present a systematic and unbiassed strategy that has enabled identification of highly immunogenic peptide epitopes derived from influenza virus and SARS-CoV-2, presented by human HLA-DR proteins. Coupling the use of HLA-DR transgenic mice with infection and vaccination with highly sensitive epitope specific cytokine ELISpot assays, we have narrowed the potential epitopes from 450-600 peptides to 5-15 peptides, by an iterative process of elimination and selection which we have termed a funnel approach. These epitopes have been validated in HLA-DR typed human CD4 T cells directly *ex vivo* and enabled derivation and implementation of HLA-DR peptide tetramers. Tetramer staining of human PBMCs enriched CD4 T memory populations from healthy adult subjects highlighting this approach as a sensitive and specific method of identifying novel epitopes and subsequent CD4 T cell responses to human viral infections.

**Importance:** Tracking single epitope-specific CD4 T cells enables sophisticated analyses of the human response to infectious pathogens, vaccines and probing the human CD4 T cell immune memory compartment. The studies presented here provide a unbiased strategy for accomplishing this goal and provide a verified compilation of candidate HLA-DR restricted CD4 T cell peptide epitopes for future studies by researchers in the field of human immunology.

## Introduction

Identification of single peptide epitopes recognized by human T cells offers tremendous advantages for understanding human cellular immunity. These epitope-specific T cells will vary in their history, number of boosts, or the original site of priming, all of which can impact their fate and effector functions. For cellular immunology studies, one notable advantage of epitope discovery is identification of single peptides that are “restricted” (e.g. bound and presented) by a known HLA molecule. Such insight can enable derivation of peptide:HLA multimers that can be used for isolation of epitope-specific T cells. Tetramer-based isolation of T cells can be followed by evaluation of T cell receptor repertoire, transcriptomics, partitioning of T cells into memory subsets, or expression of transcription factors associated with specific functions. These analyses can be performed without the need to stimulate the T cell populations, which may alter gene expression patterns (for reviews see ^1–3^). Epitope discovery and tracking of T cells with single peptide specificity can also provide insights into the factors that shape immunodominance in T cell responses, the functionality of the polyclonal responding T cells, and their persistence in the face of antigenic drift ^4–9^. Epitope discovery also enables derivation of peptide-based vaccines comprised of conserved peptide epitopes ^10–12^.

The most unbiased and comprehensive strategy for T cell epitope discovery, when possible, involves screening of elicited T cells with single peptides from libraries representing the entire translated sequence of target proteins. This approach is often infeasible because of the large volumes of PBMC needed and is complicated by the complexity of HLA molecules expressed in typical human subjects ^13^. This strategy also presents significant obstacles for pathogens expressing multiple proteins, such as bacteria, fungi, and large genome viruses. In past studies, we have used an alternative strategy for discovery of influenza CD4 T cell epitopes, using CD4 T cells from HLA-DR transgenic mice analyzed in response to vaccination or infection^6,14,15^. For large viral proteins, we implement a strategy pioneered by Tobery and colleagues using peptide matrices ^16,17^, with overlapping peptide libraries and direct *ex vivo* ELISpot assays. Using this approach, our laboratory has successfully identified HLA-DR-restricted epitopes from both seasonal and avian influenza virus proteins ^6,15^. Others have utilized a strategy termed tetramer-guided epitope mapping ^18^, involving derivation of tetramers with candidate peptides, pooling them and identification of positive tetramers within human samples, through a sequential process of de-convolution. Also, advances in machine learning have been useful for epitope discovery, particularly for HLA-class I and -class II molecules with well-defined “motifs” that correlate with peptide acquisition ^17,19–21^. Finally, a labor-intensive, technically complex, but excellent strategy for epitope discovery involves isolation of HLA-class II:peptide complex molecules from transfected cells expressing a single HLA molecule that have been either infected or pulsed with antigens, followed by elution and sequencing of the eluted, HLA-bound peptides, yielding the “immunopeptidome” of that pair of HLA-class II molecules and pathogens ^22–28^. This approach ensures that the identified peptides have a stable interaction with their presenting HLA molecules, a factor identified to be associated with immunodominance ^29,30^. This method is particularly useful for identification of peptides presented by rare HLA molecules whose peptide binding “motifs” are poorly defined (see ^31^).

For human epitope discovery from pathogens expressing multiple proteins, such as SARS-CoV-2 and influenza viruses, an efficient and informative experimental path starts with direct *ex vivo* T cell sampling after infection or vaccination of HLA-transgenic animals. This approach has the advantage of reflecting both the biochemistry of generation of HLA:peptide ligands within naturally occurring antigen presenting cells ^32–36^ and the T cell repertoire ^37–39^. Here, we describe the sequential path of empirical epitope discovery, using infected or vaccinated HLA-DR transgenic mice, in combination with cytokine ELISpot assays and overlapping peptide libraries encompassing the complete amino acid sequence of influenza virus and SARS-CoV-2 proteins to identify single viral peptide epitopes. We have followed this discovery stage by confirmation of immunogenicity in HLA-typed human PBMC and finally, have derived HLA-class II multimers on HLA-typed human samples to dissect the phenotype of circulating CD4 T cell memory cells specific for these viral pathogens.

## Materials and Methods

### Mice

The HLA-DR1 (B10.M/J-TgN-DR1) and HLA-DR4 (C57BL/6Tac-Abb<tm>TgNDR4) transgenic mice ^40,41^ were obtained from D. Zaller (Merck) through Taconic Laboratories and were maintained in the specific-pathogen-free facility at the University of Rochester according to institutional guidelines. The HLA-DR15 mice ^42^ were shared by A. Vandenbark (Oregon Health Sciences Center). HLA-DR3 mice ^43^ on the NOD background were purchased from Jackson labs (Strain #030434) and backcrossed to the class II null B6 mice developed by Mathis and colleagues ^44^. Mice were used at 2 to 8 months of age.

### Ethics statement

Mice were maintained in a specific-pathogen-free facility at the University of Rochester Medical Center according to institutional guidelines. All animal protocols adhere to the AAALAC International, the Animal Welfare Act, and the PHS guide, and were approved by the University of Rochester Committee on Animal Resources, Animal Welfare Assurance no. D16**-**00188 (A3291-01). The protocol under which these studies were conducted was originally approved 4 March 2006 (protocol 2006-030) with the most recent review and approval on 21 December 2023.

### Proteins and Peptides

Influenza peptide arrays are described in **Supplemental Table I**. Peptides were reconstituted at 10 mM in phosphate-buffered saline (PBS), with or without added dimethyl sulfoxide to increase solubility of hydrophobic peptides and 1 mM dithiothreitol for cysteine-containing peptides. Concentrated stocks (1 mM) were filter sterilized and stored at −20°C. Peptides were pooled in an array to represent each protein, or divided into multiple pools, such as HA where two pools to assess the amino and carboxy terminal portions of the protein separately were made. Spike was divided into three pools, representing the subdomains (S1-RBD, RBD and S2) to probe the different segments of the spike. Additionally, for some studies, peptides were combined into pools containing 8 to 12 peptides, where no overlapping peptides were included within any pool. These pools are arrayed into rows (labeled “R”) and columns (labeled “C”), as described previously ^16,45^ and illustrated in **Figure 1**. This strategy allows efficient identification of immunodominant peptides and exclusion of non-stimulatory peptide pools. Peptide pools were considered positive if they were at least 3-fold over background in two experiments (typically greater than 50 spots/million). Peptides in non-stimulatory pools were eliminated from further study. The sequences of the peptides used to screen were generally greater than 95% conserved with the immunizing recombinant viral proteins, depending on the availability of the overlapping peptide libraries.

**Figure 1.**
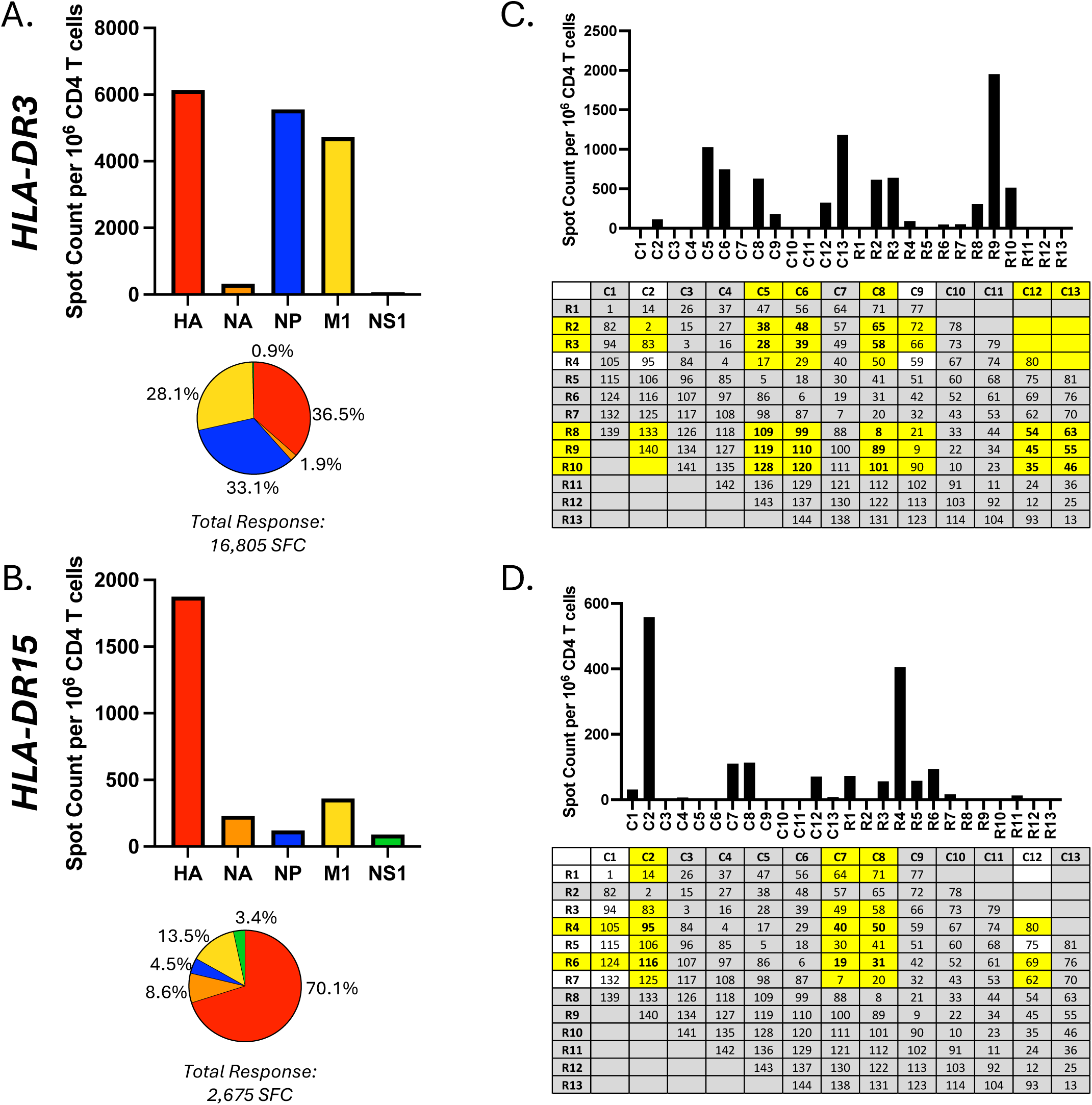
The path to identifying influenza B specific CD4 T cell epitopes. HLA-DR3 and HLA-DR15 mice were infected with influenza B/Brisbane/60/08 virus. Draining lymph nodes were harvested 12 days post infection and used as the source of CD4 T cells in IFN-γ ELISpot assays. Enriched CD4 T cells were restimulated with pools of peptides encompassing the entire translated regions of HA (red), NA (orange), NP (blue), M1 (yellow) and NS1 (green) to assess the total response to infection and allow exclusion of non-stimulatory proteins from further evaluation. Shown in A and B are the frequencies of IFN-γ producing cells per million CD4 T cells as a bar graphs, with the percent of the response to each protein shown as a pie chart with the total number of spot forming cells (SFC) shown beneath each pie. Panels C (HLA-DR3) and D (HLA-DR15) illustrate the responses to the HA peptide matrix, where peptides are grouped into pools denoted as rows (“R”) and columns (“C”) with no overlapping peptides contained in a row or column. The peptide composition of each pool is indicated with each of the 144 overlapping peptides represented by peptide numbers 1-144. Pools were considered positive with a response at least 3-fold over background (typically >50 spots). Non-stimulatory pools are indicated in gray, stimulatory pools are indicated in yellow and pools in white were stimulatory, but weak responses. Peptides at the intersection points are potentially the immunodominant epitopes, which were further tested in single peptide analyses.

### Influenza virus infections

Mice were anesthetized by intra-peritoneal injection with tribromoethanol (Avertin, 14 μL/mg body weight) and virus, diluted in 30 μL of PBS, was instilled intranasally. Influenza B/Brisbane/60/2008 and B/Florida/04/2006, provided by Richard Webby and have been previously described ^46^, A/Hawaii/70/2019 (H1N1) is a non-mouse adapted H1N1 virus, and mouse-adapted A/Switzerland/9715293/2013 (H3N2) ^47^ was kindly provided by F. Krammer. Non-lethal but immunogenic doses were determined for each virus and HLA-DR transgenic strain, typically the range for IBV was 750-1200 pfu, for H1N1 was 1000-1200 pfu and for H3N2 was 1200-2000 pfu. At 10-12 days post-infection, mice were euthanized, and single cell suspensions were prepared from mediastinal lymph nodes (mLN) and spleen. CD4 T cells were isolated by MACS (Miltenyi Biotec) negative selection following manufacturer recommendations. Purified cells were used in ELISpot assays, previously described ^15,48^.

### Protein Immunizations

HLA-DR transgenic mice were immunized subcutaneously in the rear footpads with 2-5 μg of protein in sterile PBS emulsified in Sigma Adjuvant system (Sigma) or alum (Invivogen). Recombinant SARS-CoV-2 NSP1, NSP5, NSP7/8, NSP9 and NSP15 proteins were cloned in house (Argonne National Laboratory), SARS-CoV-2 helicase protein was purchased from Sino Biological, and SARS-CoV-2 full length spike protein (Wuhan Hu-1) and nucleocapsid proteins were produced as described, respectively ^49,50^. Ten to twelve days post-vaccination mice were euthanized, and single cell suspensions were prepared from draining popliteal lymph node (pLN) and spleen and. CD4 T cells were isolated by MACS (Miltenyi Biotec) negative selection, following manufacturer recommendations and subsequently used in ELISpot assays.

### Mouse ELISpot Assays

ELISpot assays were performed as previously described ^6,15,48^. Briefly, 96-well filter plates (Millipore) were coated with 2 μg/ml purified rat anti-mouse interleukin-2 (IL-2) or IFN-γ (clone JES6-1A12 or AN-18, respectively, BD Biosciences) in PBS overnight at 4°C, washed to remove unbound antibody, and incubated with complete medium containing 10% FBS (Gibco) for 1 h to block nonspecific binding. Purified CD4 T cells from draining lymph nodes (150,000-200,000 cells per well), or spleen (300,000 cells per well) were co-cultured with 500,000 syngeneic splenocytes (HLA-DR4, HLA-DR3, or HLA-DR15). For DR1-Tg mice, that express the mouse I-A^f^ molecule, which poorly selects a CD4 T cell repertoire, DAP.3 fibroblast cells transfected with the genes encoding HLA-DR1, generously provided by E. Long, (NIAID, NIH; 35,000 cells/well) were used, with un-transfected DAP.3 cells used as a control. Cultures were incubated with either a pool of peptides or a single peptide at a final concentration of 0.5 μM or 1μM, respectively, in a total volume of 200 μl for 18 to 20 h at 37°C and 5% CO_2_. Plates were washed (1× PBS, 0.1% Tween20) and biotinylated rat anti-mouse IL-2 or IFN-γ (clone JES6-5H or XMG1.2, respectively, BD Biosciences) was added (2 μg/ml in wash buffer with 10% FBS) and incubated at room temperature for 30 min. After washing, streptavidin-conjugated alkaline phosphatase (JacksonImmuno Research) was added at a dilution of 1:1000 (1mg/ml stock) in wash buffer with 10% FBS at 50 μl/well and incubated for 30 min at room temperature, washed and developed using Vector Blue substrate (Vector Laboratories) prepared in 100 mM Tris (pH 8.2). Quantification of spots was performed with an Immunospot reader series 5.2. CD4 T cells, APC and media with no added peptide were used for negative control, background responses, and all conditions were performed in duplicate or triplicate. All responses with background subtracted are shown, average values that are at least 3-fold over background were considered a positive response.

### Human ELISpot Assay

HLA-typed human PBMC samples were obtained from C.T.L. (Shaker Hts, OH). Samples were collected from health adults aged 19-54 between 2018 and 2023 and stored in the vapor phase of LN_2_ until time of use. After thawing at 37°C and overnight rest in culture, cells were depleted of CD8 and CD56 cells using MACS microbeads per manufacturer instructions (Miltenyi Biotec). ELISpot assays were performed as previously described ^51^. Briefly, CD8- and CD56-depleted PBMCs (300,000-400,000 cells per well) were cultured with single peptides on plates coated with 10 μg/ml anti-human IFNγ (clone 1-D1K, MabTech) for 36 hr at 37°C, 5% CO_2_. After incubation, plates were washed and incubated with biotinylated detection antibody human IFNγ (2 μg/ml, clone 7-B6-1, MabTech) for 2 h followed by washing and incubation for 30min with streptavidin conjugated-alkaline phosphatase (1:1000 dilution of 1mg/ml stock) and development using Vector Blue substrate. Quantification of cytokine-secreting cells was performed as described above. Data are presented as the frequency of cytokine-producing cells per million CD8- and CD56-depleted PBMCs with background response subtracted. All responses with background subtracted are shown, including zero values. Median responses three-fold over background (average background 5 spots) are considered positive responses.

### Human Donor HLA Typing for tetramer analyses

HLA typing of aphaeresis donors for tetramer analyses was performed using the AllType NGS 11-Loci Amplification Kit (One Lambda) according to manufacturer’s instructions. Resulting libraries were sequenced on MiSeq lane at 150×150bp. HLA types were called using the TypeStream Visual Software from One Lambda. Screening was first performed by ELISpot assay.

### Preparation of U-Load peptide-HLA monomers and tetramer assembly

Unfolded, biotinylated U-Load monomers (Immudex) were obtained for HLA-DRB1*04:01. Influenza virus peptide IAV M1 p17 was commercially synthesized, diluted to 1 mM in DMSO, and loaded into U-Load HLA-DRB1*04:01 according to the manufacturer’s instructions at 37°C for 20h. An HLA-DRB1*04:01 negative control, containing the CLIP peptide, was also folded. Custom HLA-DRB1*04:01 monomers, loaded with IAV H3p81 or SARS-CoV-2 Sp8, were generated by Immudex. Peptide-HLA tetramers were then assembled according to the manufacturer’s instructions. Briefly, 0.48ng of fluorochrome-conjugated streptavidin (PE or PE-Cy7, Biolegend) was added to peptide-HLA complex (500 nM in 1x PBS and 5% v/v glycerol) in three equal volumes for a total of 60 μL. After each 1/3 volume addition samples were mixed and incubated for 10 min at 4°C in the dark. Assembled tetramers were stored at 4°C in the dark until use.

### Tetramer-associated magnetic enrichment

PBMCs were isolated from apheresis rings obtained from the St. Jude Children’s Research Hospital Blood Donor Center under Department of Pathology protocol BDC035. All apheresis rings were de-identified before release. PBMCs from donors with confirmed HLA-DRB1*04:01 were selected. 3-12×10^6^ PBMCs were blocked with human TruStain FcX blocking buffer (Biolegend) for 15 min at 4°C in FACS Buffer (1x PBS, 0.5% BSA, 0.5 mM EDTA) followed by centrifugation at 500 xg for 5 min and re-suspension in 50 nM dasatinib (Sigma-Aldrich) in 1x PBS and incubated for 30 min at 37°C and 5% CO_2_. PBMCs were then centrifuged at 500 xg for 5 min and stained with pairs of PE- and PE/Cy7-conjugated tetramers (1.5 μL tetramer per 1×10^6^ PBMCs) for each target (1:10 dilution in FACS buffer containing 500 μM D-biotin) in a volume 100 μL. PBMCs were incubated with tetramers at 25°C for 1 h with gentle mixing every 15 min. Tetramer-associated magnetic enrichment (TAME), using Miltenyi LS columns, was used to select tetramer-bound cells as per the manufacturer’s instructions. Briefly, tetramer-stained PBMCs were washed in 1 mL cold MACS buffer (1x PBS, 0.5% BSA, 2 mM EDTA) then suspended in 80 μL cold MACS buffer. 20 μL each of anti-PE and anti-Cy7 microbeads (Miltenyi) were added, cells were gently mixed, and incubated on ice for 30 min. Cells were then washed in 1 mL cold MACS buffer and re-suspended in 500 μL MACS buffer before adding to an LS column. The flow-through cells were collected. The columns were removed from the magnet and washed to collect the tetramer-bound (TAME) fraction. The TAME cell fractions as well as an aliquot of PBMCs not run through the column were pelleted and suspended in FACS buffer containing 500 μM D-Biotin (FACS-B) and a cocktail of fluorophore-conjugated surface antibodies (**Supplemental Table II**) in a final volume of 100 μL and incubated in the dark on ice for 30 min. PBMCs were then washed twice and suspended in 250 μL FACS-B. Samples were analyzed on a 5-laser Aurora spectral cytometer. Cell population gating and fluorescence analysis was performed using FlowJo version 10.7.2 software (BD Biosciences). Additional analysis was performed in R (v4.1.0).

## Results

A key strategy for these studies was to use humanized mice, expressing HLA-DR molecules, to identify pathogen-derived peptides recognized by epitope-specific CD4 T cells as the first step for ultimate validation in human CD4 T cells. The approach focused on four different and commonly expressed HLA-DR alleles: HLA-DR1 (Β1:0101), HLA-DR4 (Β1:0401), HLA-DR15 (Β1:1501), and HLA-DR3 (Β1:0301), for which CD4 T cell epitope discovery was enabled by acquisition of HLA-DR transgenic mice. Epitope discovery was initiated with either infection or vaccination of mice (depending on the virus and availability of recombinant proteins for vaccination), where the isolated CD4 T cells were tested directly *ex vivo* at the peak of the primary response. Shown in **Figure 1** are examples of the experimental path of epitope discovery for infection of HLA-DR3 and HLA-DR15 mice with influenza B virus (IBV). In many cases, the focus of the CD4 T cell response was initially assessed by quantifying cytokine production to overlapping peptide pools encompassing the entire translated sequence of the individual candidate proteins (HA, NA, M1, NP and NS1), using cytokine ELISpots. Using this approach, we found that for HLA-DR3 (**Figure 1A**), the response focused equally on HA, NP and M1, with minimal responses to NA and NS1. These negative responses enabled elimination of these proteins from further analyses. In contrast, the response in HLA-DR15 mice to IBV (**Figure 1B**) was dominated by HA, representing almost 75% of the total response, with negligible responses specific for NP and NS1 and only a minor response to M1. Based on these results, NA, NP and NS1 were eliminated from consideration and single peptide epitopes were defined for the dominant influenza virus proteins in both strains of mice. **Figure 1C** illustrates the results from a peptide pooling matrix strategy for HA-B for HLA-DR3 and HLA-DR15 (**Figure 1D**), where individual peptides are pooled in such a way that no adjacent peptides are in the same small pool of peptides^16^. This strategy allows elimination of many peptides from further consideration (in grey) and identification of peptides in the most stimulatory pools (in yellow). The peptide candidates were tested in subsequent experiments shown in **Figure 2**, allowing identification of several dominant peptides (e.g. HA p55) with several minor peptides restricted to HLA-DR3 **(Figure 2A)** and one major HA peptide restricted to HLA-DR15 (HA p95) **(Figure 2B)**, with one or more minor epitopes. For smaller proteins, such as M1, single peptides were individually tested. Single peptides were tested and confirmed in at least 2 experiments. **Figure 2C and 2D** show the results of single peptide experiments for M1 for HLA-DR3 and HLA-DR15, where each of the peptides in the peptide array was tested. Epitope discovery was continued using these strategies for IAV. The summary of the epitope discovery for influenza virus responses is shown in **Figure 3** and listed in **Tables I-IV**, for each of the mouse strains, that indicate both the peptide number described in each array and the starting and ending amino acid, the sequence of each peptide, and the method by which epitopes were mapped.

**Figure 2.**
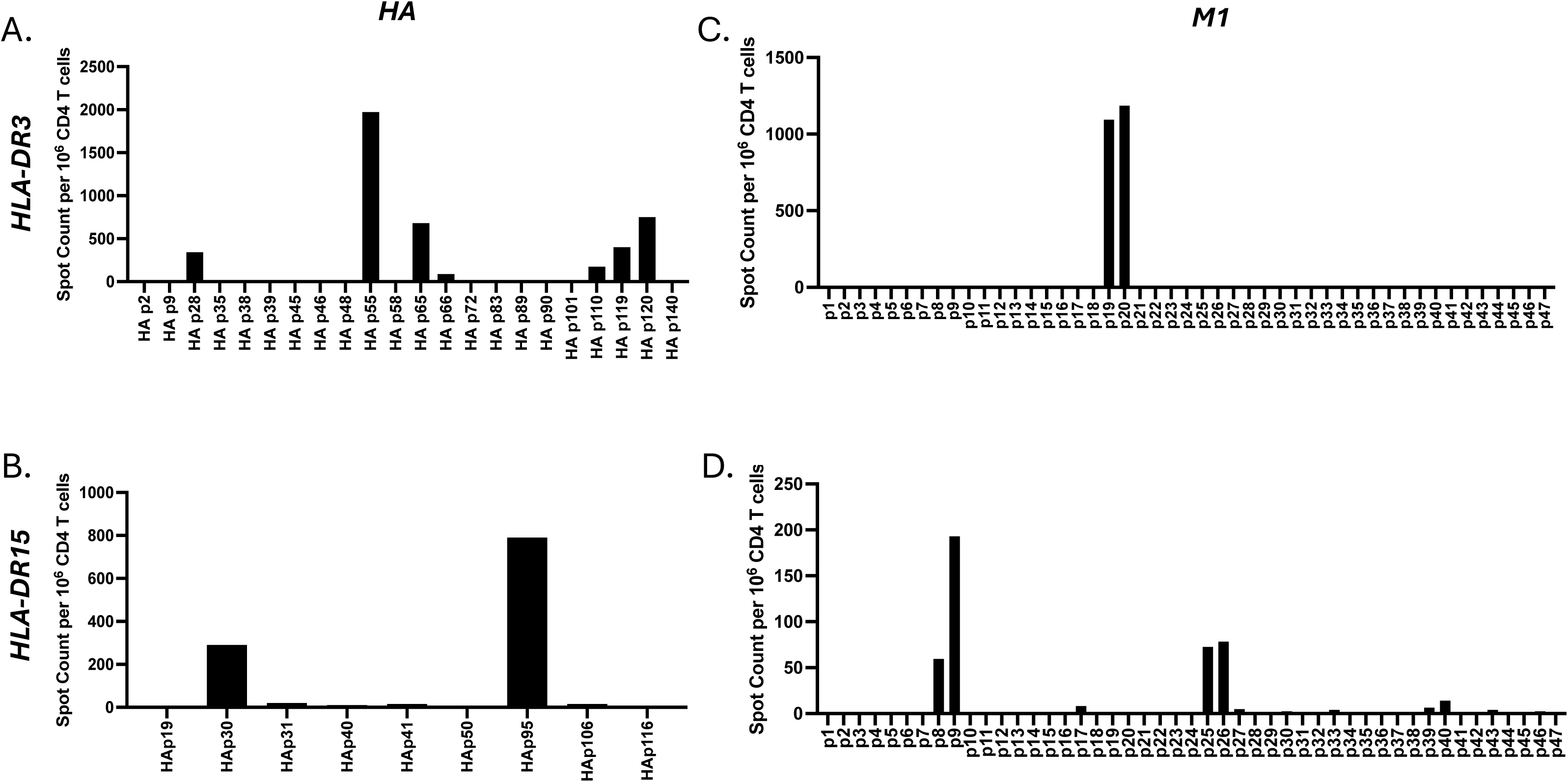
Single peptide analysis of the HA and M1 proteins of influenza B-specific CD4 T cell epitopes detected after infection in HLA-DR3 and HLA-DR15 transgenic mice. Shown here are representative plots of single peptide analyses for identification of CD4 T cell peptide epitopes in HA (left) and M1 (right) proteins of influenza B/Brisbane/60/08 after infection. The peptides selected for analysis for HA-B are based on the stimulatory peptides and peptides eliminated in the matrices shown in Figure 1. M1 peptides were tested individually without using a matrix pooling approach. Shown are the responses of splenic CD4 T cells with the bar graphs represent the average frequency of IFN-γ producing cells per million CD4 T cells with background subtracted.

**Figure 3.**
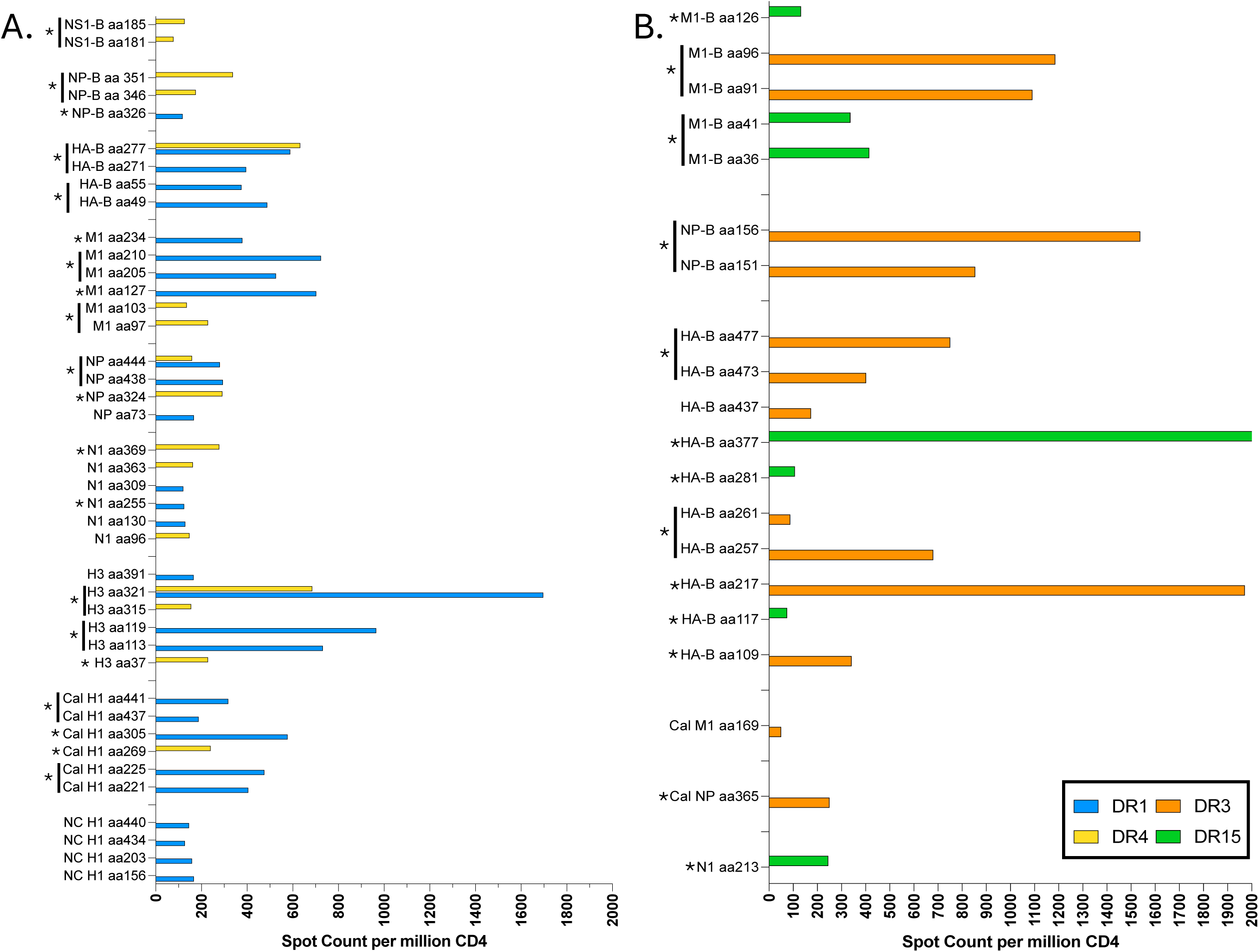
Summary of single influenza H1N1, H3N2 and influenza B peptide-epitopes identified in HLA-DR Tg mice. Shown are the average frequencies of cytokine producing cells per million CD4 cells in HLA-DR1 (blue), HLA-DR4 (yellow), HLA-DR3 (orange) and HLA-DR15 (green) transgenic mice. Epitopes that were tested in healthy adult human PBMC samples, shown in Figure 9 are indicated with an asterisk. Adjacent overlapping peptides were tested as a combined peptide and are indicated by a line next to the asterisk.

**Table I.**
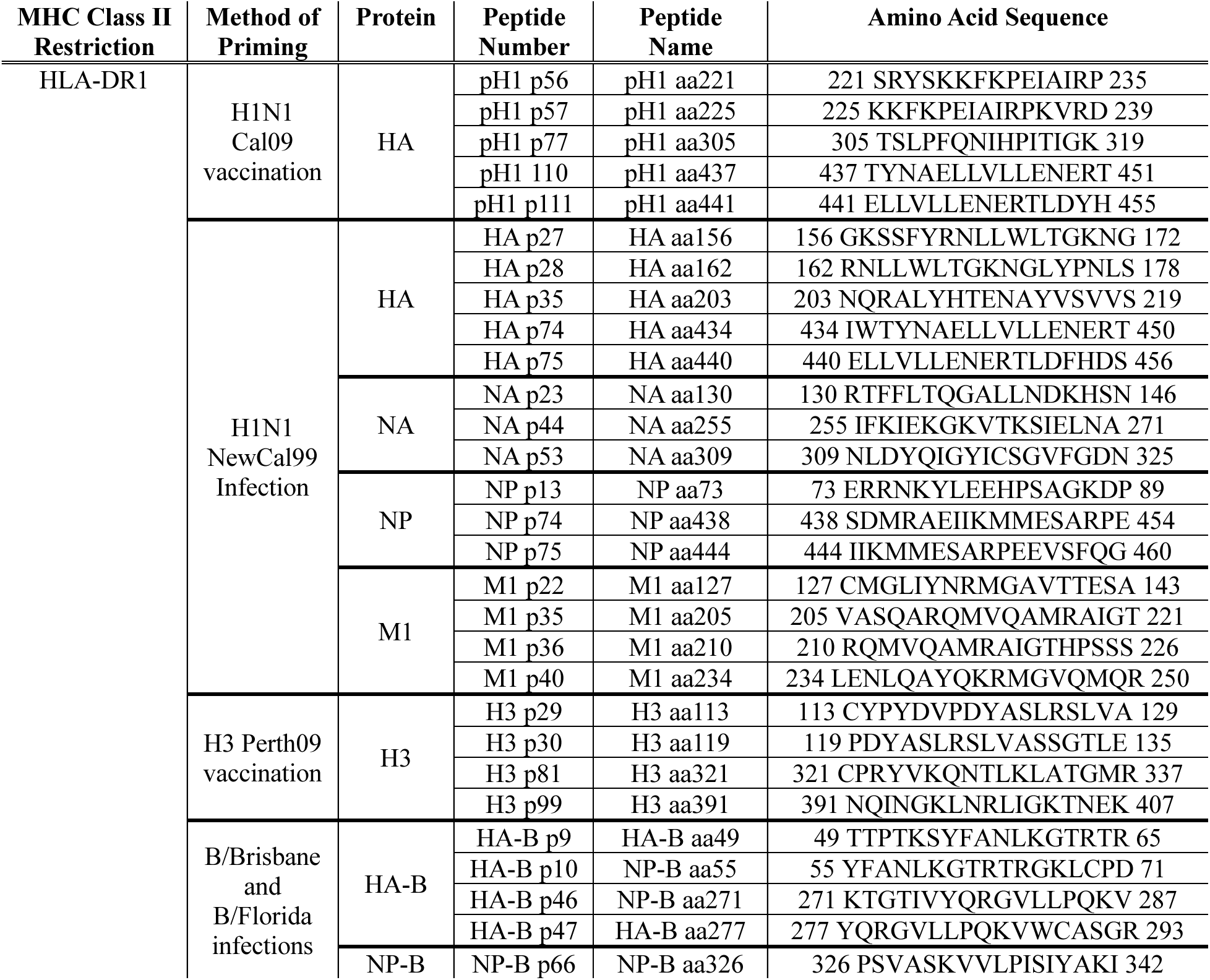
HLA-DR1 peptide sequences with amino acid numbers, peptide names and numbers and the method of priming used to map the peptide epitopes in influenza A H1N1 and influenza B.

**Table II.** HLA-DR4 peptide sequences with amino acid numbers, peptide names and numbers and the method of priming used to map the peptide epitopes in influenza A H1N1 and influenza B.

| MHC Class II Restriction | Method of Priming | Protein | Peptide Number | Peptide Name | Amino Acid Sequence |
| --- | --- | --- | --- | --- | --- |
| HLA-DR4 | H1N1 Cal09 vaccination | HA | HA p68 | HA aa269 | 269 RYAFAMERNAGSGII 283 |
|  | H1N1 NewCal99 Infection | NA | NA p17 | NA aa96 | 96 GWAITYTKDNSIRIGSKG 112 |
|  |  |  | NA p62 | NA aa363 | 363 KSNRLRKGFEMIWDPNG 379 |
|  |  |  | NA p63 | NA aa369 | 369 KGFEMIWDPNGWTDTS 385 |
|  |  | NP | NP p55 | NP aa324 | 324 HKSQVLVWMA CNSAAFED 340 |
|  |  |  | NP p75 | NP aa444 | 444 IIKMMESARPEEV SFQG 460 |
|  |  | M1 | M1 p17 | M1 aa97 | 97 VKLYRKLKREITFHGAK 113 |
|  |  |  | M1 p18 | M1 aa103 | 103 LKREITFHGAK EIALSY 119 |
|  | H3 Perth09 vaccination | H3 | H3 p10 | H3 aa37 | 37 PNGTIVKTITNDQIEVT 53 |
|  |  |  | H3 p80 | H3aa315 | 315 RITYGACPRYVKQNTLK 331 |
|  |  |  | H3 p81 | H3 aa321 | 321 CPRYVKQNTLKLATGMR 337 |
|  | B/Brisbane and B/Florida infections | HA-B | HA-B p47 | HA-B aa277 | 277 YQRGVLLPQKVWCASGR 293 |
|  |  | NP-B | NP-B p70 | NP-B aa346 | 346 GFNVEEYS MVGYEAMAL 362 |
|  |  |  | NP-B p71 | NP-B aa351 | 351 EYS MVGYEAMALLYNMAT 367 |
|  |  | NS1-B | NS1-B p46 | NS1-B aa181 | 181 NVLSLRVLVNGTFLKH 196 |
|  |  |  | NS1-B p47 | NS1-B aa185 | 185 LRVLVNGTFLKHPNGY 200 |

**Table III.** HLA-DR3 peptide sequences with amino acid numbers, peptide names and numbers and the method of priming used to map the peptide epitopes in influenza A H1N1 and influenza B.

| MHC Class II Restriction | Method of Priming | Protein | Peptide Number | Peptide Name | Amino Acid Sequence |
| --- | --- | --- | --- | --- | --- |
| HLA-DR3 | H1N1 infection | NP | NP p92 | NP aa365 | 365 IASNENVETMDSNTL 379 |
|  | B/Brisbane infection | HA-B Brisbane | HA-B p28 | HA-B aa109 | 109 CFPIMHDRTKIRQLP 123 |
|  |  |  | HA-B p55 | HA-B aa217 | 217 AKLYGDSKPQKFTSS 231 |
|  |  |  | HA-B p65 | HA-B aa257 | 257 QSGRIVVDY MVQKSG 271 |
|  |  |  | HA-B p66 | HA-B aa261 | 261 IVVDY MVQKSGKTGT 275 |
|  |  |  | HA-B p110 | HA-B aa437 | 437 NEILELDEKVDDLRA 451 |
|  |  |  | HA-B p119 | HA-B aa473 | 473 EDEHLLALERKLKKM 487 |
|  |  |  | HA-B p120 | HA-B aa477 | 477 LLALERKLKMLGPS 491 |
|  |  | NP-B | NP-B p31 | NP-B aa151 | 151 GGTFYKMVRDDKTIYFS 167 |
|  |  |  | NP-B p32 | NP-B aa156 | 156 KMVRDDKTIYFSPIRIT 172 |
|  |  | M1-B | M1-Bp19 | M1-B aa91 | 91 TKKKGLILAERKMRRVCV 107 |
|  |  |  | M1-Bp20 | M1-B aa96 | 96 LILAERKMRRCVSFHEA 112 |

**Table IV.** HLA-DR15 peptide sequences with amino acid numbers, peptide names and numbers and the method of priming used to map the peptide epitopes in influenza A H1N1 and influenza B.

| MHC Class II Restriction | Method of Priming | Protein | Peptide Number | Peptide Name | Amino Acid Sequence |
| --- | --- | --- | --- | --- | --- |
| HLA-DR15 | H1N1 infection | NA | NA p54 | NA aa213 | 213 TDTIKSWRNNILRTQ 227 |
|  | B/Brisbane infection | HA-B Brisbane | HA-B p30 | HA-B aa117 | 117 TKIRQLPNLLRGYEH 131 |
|  |  |  | HA-Bp71 | HA-B aa281 | 281 GILLPQKVWCASGRS 295 |
|  |  |  | HA-Bp95 | HA-B aa377 | 377 EGMIAGWHGYTSHGA 391 |
|  |  | M1-B | M1-Bp8 | M1-B aa36 | 36 EFDLDSALEWIKNKRCL 52 |
|  |  |  | M1-Bp9 | M1-B aa41 | 41 SALEWIKNKRCLTDIQK 57 |
|  |  |  | M1-Bp26 | M1-B aa126 | 126 YCLMVMYLNPGNYSMQV 142 |

To extend epitope discovery to SARS-CoV-2, recombinant protein vaccination was used, followed as before, by direct *ex vivo* ELISpot analyses. For SARS-CoV-2 spike, a large protein consisting of over 250 overlapping peptides, the peptide libraries were separated into three domains for initial screening. The segments and boundaries across spike are shown in **Figure 4A**. The receptor binding domain (“RBD”) consisted of 47 overlapping peptides, the S1 segment minus the RBD (“S1-RBD”), consisted of 87 peptides, and “S2”, which is the entire conserved membrane proximal domain consisted of 119 overlapping peptides. Peptides at the boundaries of each segment were included in both pools. **Figures 4B and 4C** show the results of testing of each of the three segments after the HLA-DR transgenic mouse strains were vaccinated with recombinant full-length SARS-CoV-2 spike protein, formulated with adjuvant. The dominance pattern observed was unique to each strain, indicating strong HLA-DR-linked epitope selection. For example, DR1-restricted responses were dominated by RBD and S1-RBD-derived peptides, with only minor responses to S2, whereas DR3 and was dominated by the smaller RBD and S2-derived peptide pools. These results were followed by the peptide pooling matrix analyses **(Figure 5)**. The elimination/selection experiments were followed by single peptides analyses, shown in **Figure 6**. This iterative process enabled selection from 253 total spike-derived peptides to 6 peptides presented by HLA-DR1, 6 from HLA-DR3, 5 from HLA-DR4 and 3 from HLA-DR15, when using a frequency of 100 cytokine producing cells per million as an “immunodominant positive” peptide. These studies were next extended to other SARS-CoV-2-derived peptides to nucleocapsid (N), helicase, and a subset of non-structural proteins (NSPs). The choices across SARS-CoV-2 proteins were in part based on their availability and in part based on homology with seasonal human coronaviruses (HCoV), where amino acid sequence alignments suggested the potential for recall from T cell memory established by these HCoV, that are prominent for helicase and N which we anticipated would enrich for reactivity in human CD4 T cells. **Figure 7A and 7B** show the results with epitope discovery in the 4 strains of HLA-DR transgenic mice for SARS-CoV-2-derived epitopes. **Tables V-VIII** also indicate all individual epitopes with peptide and amino acid numbers and sequences. It should be noted that responses in the different strains of mice exhibit characteristic relative immunogenicity in responses after infection or vaccination. These differences may relate to the site of integration of the transgene, or selection of the CD4 T cell repertoire in the host. We have no evidence that the HLA-DR transgenic mice differ in the fraction of cells that are positive for HLA-DR or the cell surface density of HLA-DR molecules.

**Figure 4.**
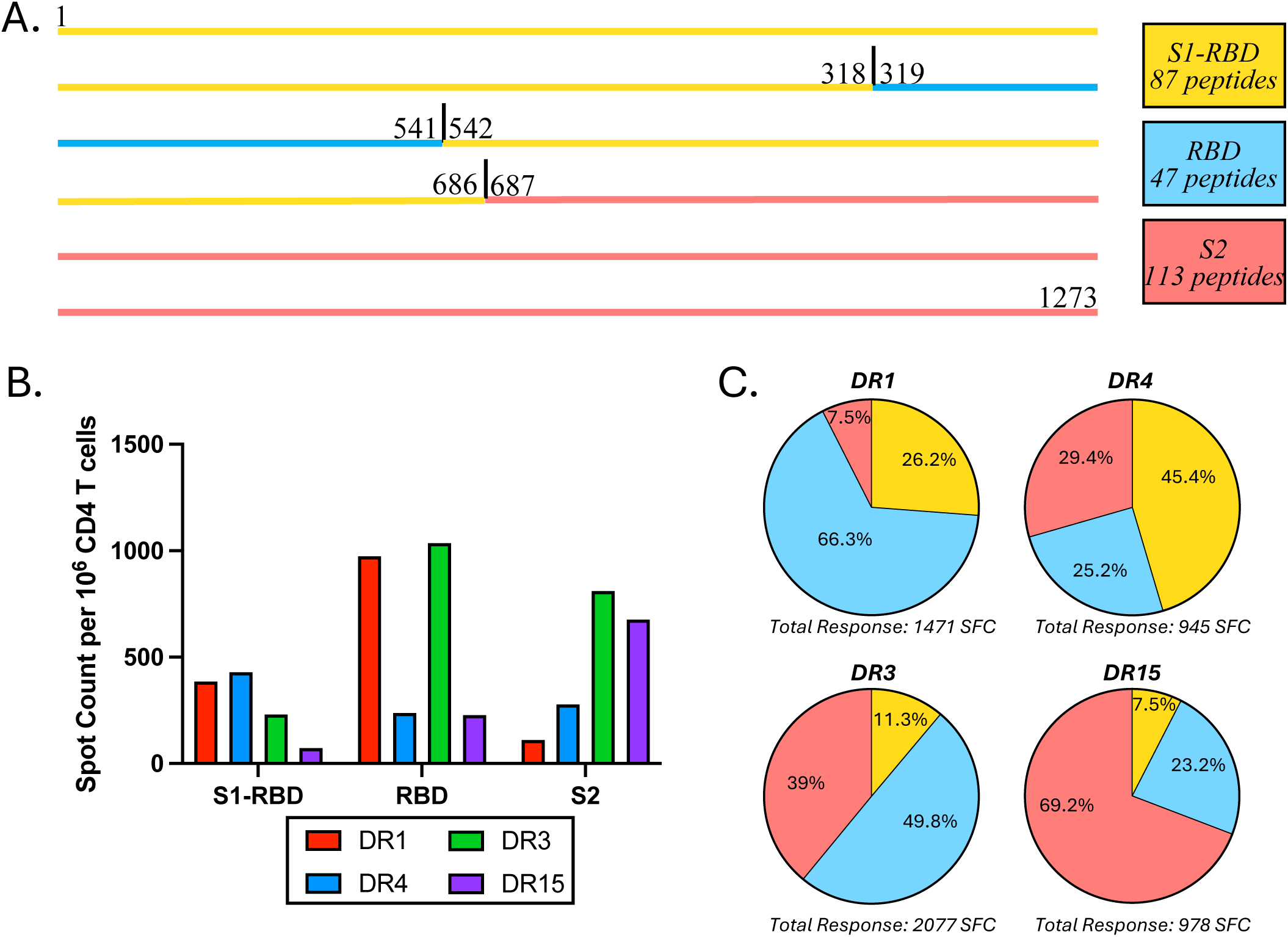
HLA-DR transgenic mice have distinct immunodominance patterns to SARS-CoV-2 spike protein. The responses to recombinant spike protein were evaluated in IL-2 ELISPOT assays following vaccination with SARS-CoV-2 Spike, using overlapping peptides representing each segment indicated in the colored boxes to the right. (B) CD4 T cell responses were evaluated by IL-2 ELISPOT assays in HLA-DR1 (red), DR4 (blue), DR3 (green) and DR15 (purple) that had been vaccinated with recombinant SARS-CoV-2 spike protein as described in Materials and Methods. Responses are shown as the frequency of IL-2 producing cells per million CD4 T cells for each of the overlapping peptide libraries indicated in A. Panel C illustrates the percent of the response to each segment of spike (S1-RBD (yellow), RBD (blue) and S2 (salmon)) and the total response to SARS-CoV-2 Spike protein indicated below each pie.

**Figure 5.**
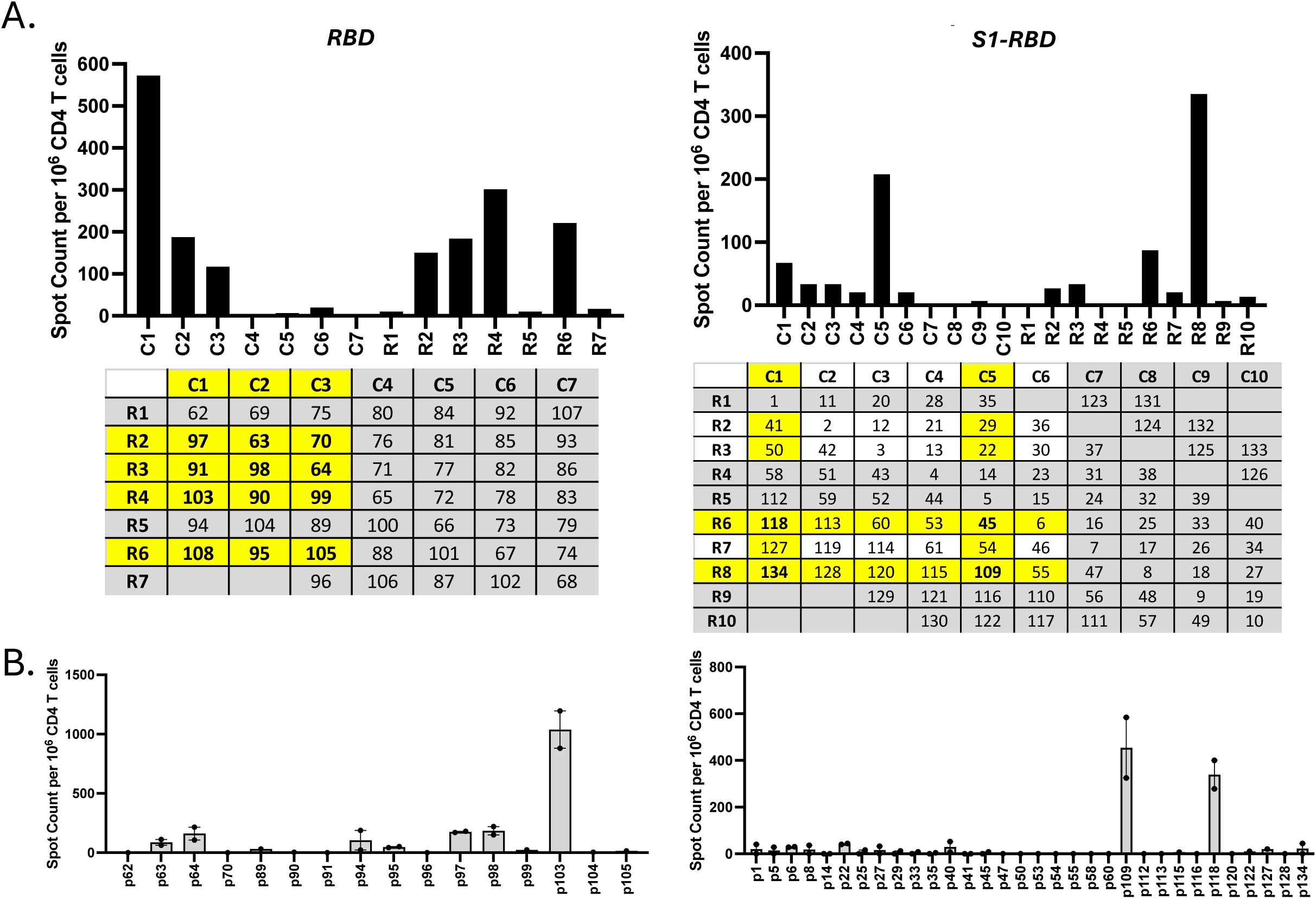
Path of mapping single peptide epitopes in SARS-CoV-2 spike protein. Example of matrices for spike in HLA-DR1 mice from the spike domains showing greatest reactivity. (RBD, left and S1-RBD, right) (see Figure 4). Candidate single peptides from the results of the matrix, highlighted in yellow, were tested in the subsequent assay shown in B.

**Figure 6.**
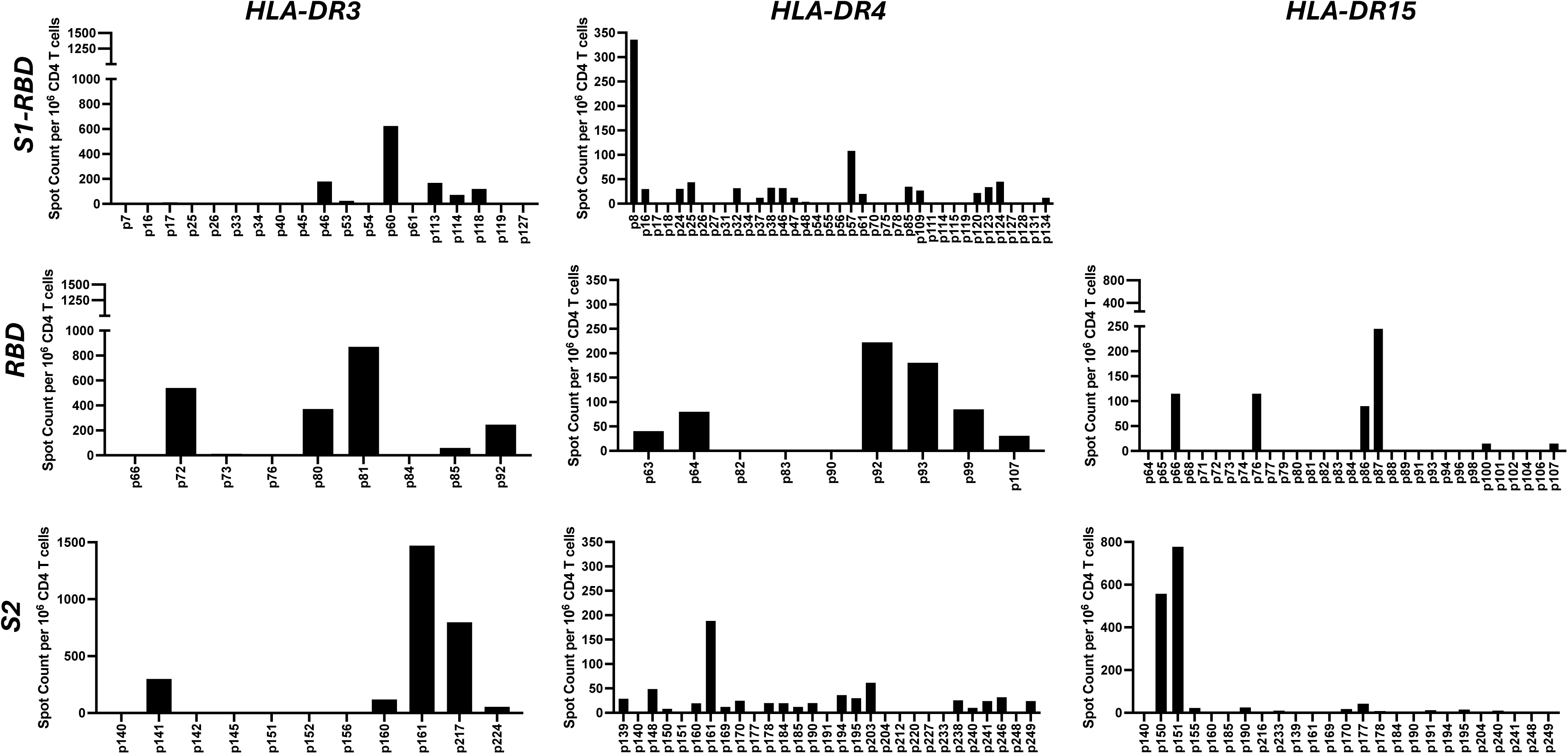
Single peptide analysis of the spike protein for HLA-DR3, HLA-DR4 and HLA-DR15. Shown is an example of single peptide analyses following vaccination with SARS-CoV-2 Spike protein, based on the results of peptide pooling matrices, leading to testing of single candidate peptides that are unique to each HLA-DR allele. There was negligible reactivity to epitopes contained in the S-RBD segment in HLA-DR15 transgenic mice and these data are not shown here.

**Figure 7.**
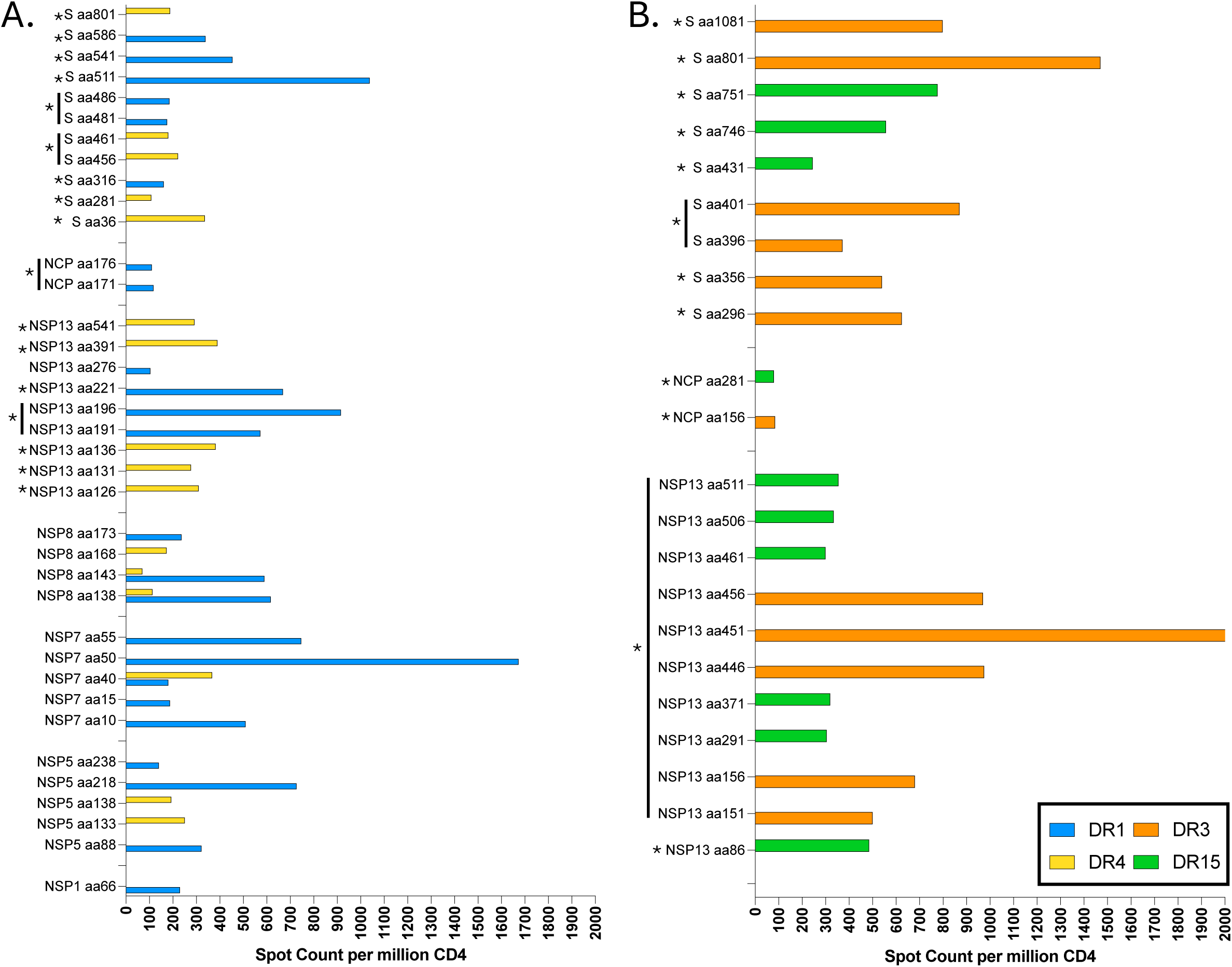
Summary of single SARS-CoV-2 CD4 T cell peptide-epitopes identified in HLA-DR transgenic mice. Shown is the average frequency of cytokine producing cells per million CD4 cells in HLA-DR1 (blue), HLA-DR4 (yellow), HLA-DR3 (orange) and HLA-DR15 (green) transgenic mice that had been vaccinated with recombinant SARS-CoV-2 proteins. Epitopes that were tested in healthy adult human PBMC samples, shown in Figure 9, are indicated with an asterisk. Adjacent overlapping peptides were tested as a combined peptide and are indicated by a line next to the asterisk. In some cases, helicase peptides were tested as a selected pool.

**Table V.** HLA-DR1 peptide sequences with amino acid numbers, peptide names and numbers used in the figures and the method of priming used to map the peptide epitopes in SARS-CoV-2.

| MHC Class II Restriction | Method of Priming | Protein | Peptide Number | Peptide Name | Amino Acid Sequence |
| --- | --- | --- | --- | --- | --- |
| HLA-DR1 | Protein vaccination | Nucleocapsid | NCP p35 | NCP aa171 | 171 FYAEGSRGGSQASSRSS 187 |
|  |  |  | NCP p36 | NCP aa176 | 176 SRGGSQASSRSSSRN 192 |
|  |  | Helicase | NSP13 p39 | NSP13 aa191 | 191 SKVQIGEYTFEKGDYGD 207 |
|  |  |  | NSP13 p40 | NSP13 aa196 | 196 GEYTFEKGDYGDVAVYR 212 |
|  |  |  | NSP13 p45 | NSP13 aa221 | 221 VGDFYFVLTSHTVMPLSA 237 |
|  |  |  | NSP13 p56 | NSP13 aa276 | 276 KYSTLQGPPGTGKSHFA 292 |
|  |  | NSP1 | NSP1 p14 | NSP1 aa66 | 66 QPYVFIKRSDARTAPHG 72 |
|  |  | NSP5 | NSP5 p21 | NSP5 aa88 | 88 LKVDTANPKTPKYKFVR 104 |
|  |  |  | NSP5 p47 | NSP5 aa218 | 218 FLNRFTTTLNDFNLVAM 234 |
|  |  |  | NSP5 p51 | NSP5 aa238 | 238 YEPLTQDHVDILGPLSA 254 |
|  |  | NSP7 | NSP7 p5 | NSP7 aa10 | 10 SVVLLSVLQQLRVESSS 26 |
|  |  |  | NSP7 p6 | NSP7 aa15 | 15 SVLQQLRVESSSKLWAQ 31 |
|  |  |  | NSP7 p11 | NSP7 aa40 | 40 LLAKDTTEAFEKMSVSL 56 |
|  |  |  | NSP7 p13 | NSP7 aa50 | 50 EKMVSLLSVLLSMQGA 66 |
|  |  |  | NSP7 p14 | NSP7 aa55 | 55 LLSVLLSMQGAVDINKL 71 |
|  |  | NSP8 | NSP8 p31 | NSP8 aa138 | 138 YKNTCDGTTFTYASALW 154 |
|  |  |  | NSP8 p32 | NSP8 aa143 | 143 DGTFTYASALWEIQV 159 |
|  |  |  | NSP8 p38 | NSP8 aa173 | 173 SMDNSPNLAWPLIVTAL 189 |
|  |  | Spike | Spk p64 | Spk aa316 | 316 SNFRVQPTESIVRFPNI 332 |
|  |  |  | Spk p97 | Spk aa481 | 481 NGVEGFNCYFPLQSYGF 497 |
|  |  |  | Spk p98 | Spk aa486 | 486 FNCYFPLQSYGFQPTNG 502 |
|  |  |  | Spk p103 | Spk aa511 | 511 VVLSFELLHAPATVCGP 527 |
|  |  |  | Spk p109 | Spk aa541 | 541 FNFNGLTGTGVLTESNK 557 |
|  |  |  | Spk p118 | Spk aa586 | 586 DITPCSFGGVSITPGT 602 |

**Table VI.**
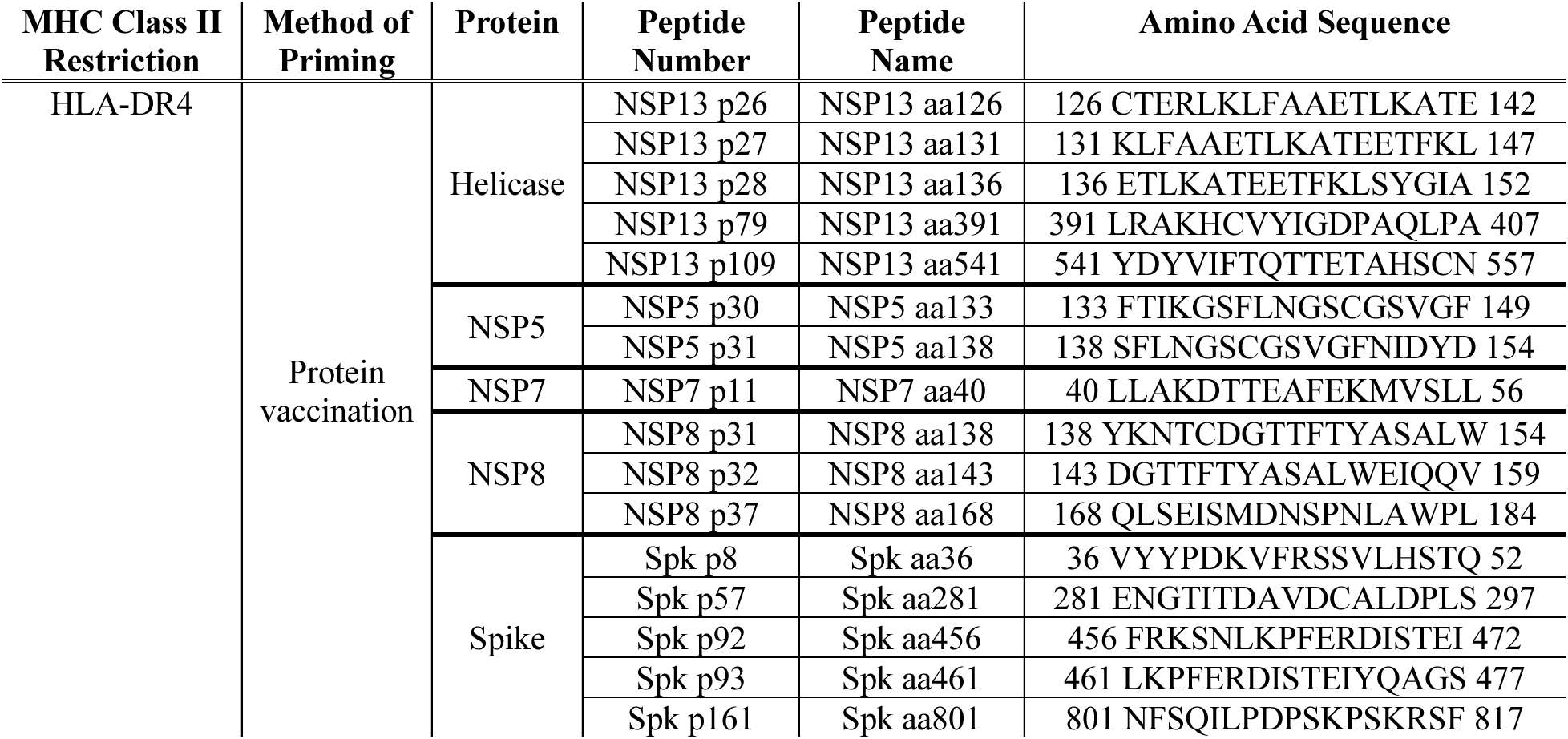
HLA-DR4 peptide sequences with amino acid numbers, peptide names and numbers used in the figures and the method of priming used to map the peptide epitopes in SARS-CoV-2.

**Table VII.**
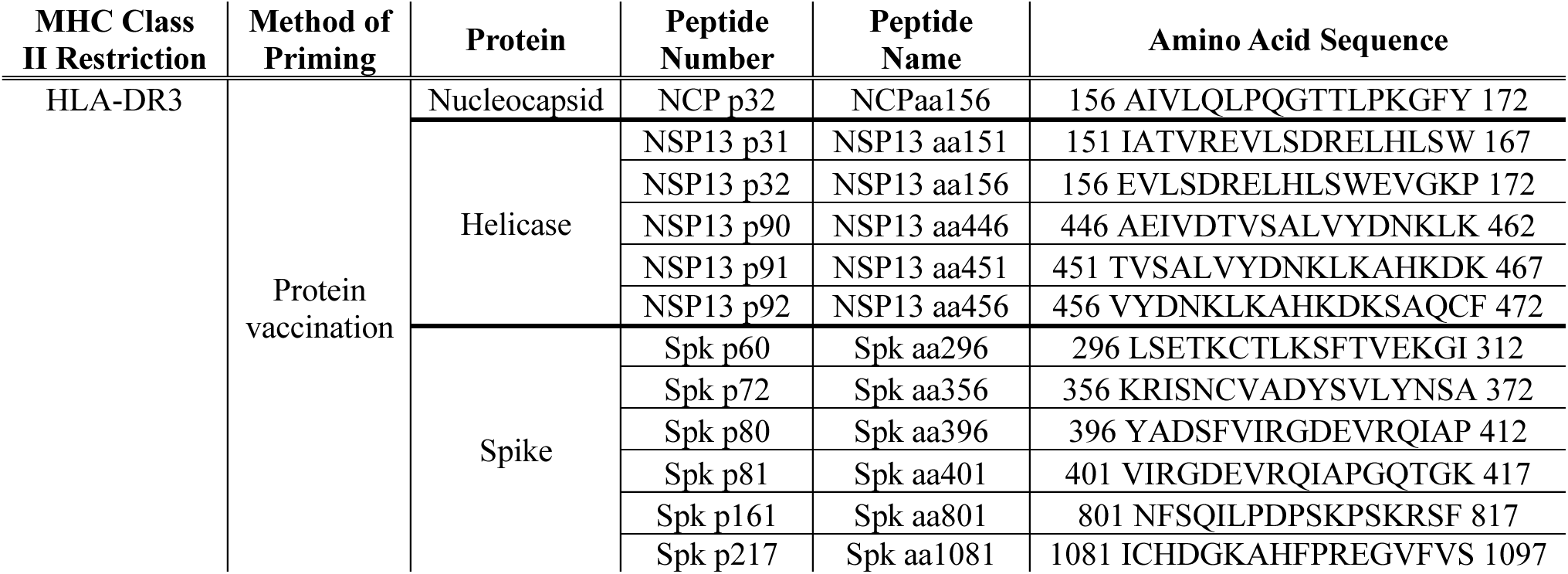
HLA-DR3 peptide sequences with amino acid numbers, peptide names and numbers used in the figures and the method of priming used to map the peptide epitopes in SARS-CoV-2.

| MHC Class II Restriction | Method of Priming | Protein | Peptide Number | Peptide Name | Amino Acid Sequence |
| --- | --- | --- | --- | --- | --- |
| HLA-DR3 | Protein vaccination | Nucleocapsid | NCP p32 | NCPaa156 | 156 AIVLQLPQGTTLPKGFY 172 |
|  |  | Helicase | NSP13 p31 | NSP13 aa151 | 151 IATVREVLSDRELHLSW 167 |
|  |  |  | NSP13 p32 | NSP13 aa156 | 156 EVLSDRELHLSWEVGKP 172 |
|  |  |  | NSP13 p90 | NSP13 aa446 | 446 AEIVDTVSALVYDNKLK 462 |
|  |  |  | NSP13 p91 | NSP13 aa451 | 451 TVSALVYDNKLKAHKDK 467 |
|  |  |  | NSP13 p92 | NSP13 aa456 | 456 VYDNKLKAHKDKSAQCF 472 |
|  |  | Spike | Spk p60 | Spk aa296 | 296 LSETKCTLKSFTVEKGI 312 |
|  |  |  | Spk p72 | Spk aa356 | 356 KRISNCVADYSVLYNSA 372 |
|  |  |  | Spk p80 | Spk aa396 | 396 YADSFVIRGDEVQRQIAP 412 |
|  |  |  | Spk p81 | Spk aa401 | 401 VIRGDEVQRQIAPGQTGK 417 |
|  |  |  | Spk p161 | Spk aa801 | 801 NFSQILPDPSKPSKRSF 817 |
|  |  |  | Spk p217 | Spk aa1081 | 1081 ICHDGKAHFPREGVFVS 1097 |

**Table VIII.**
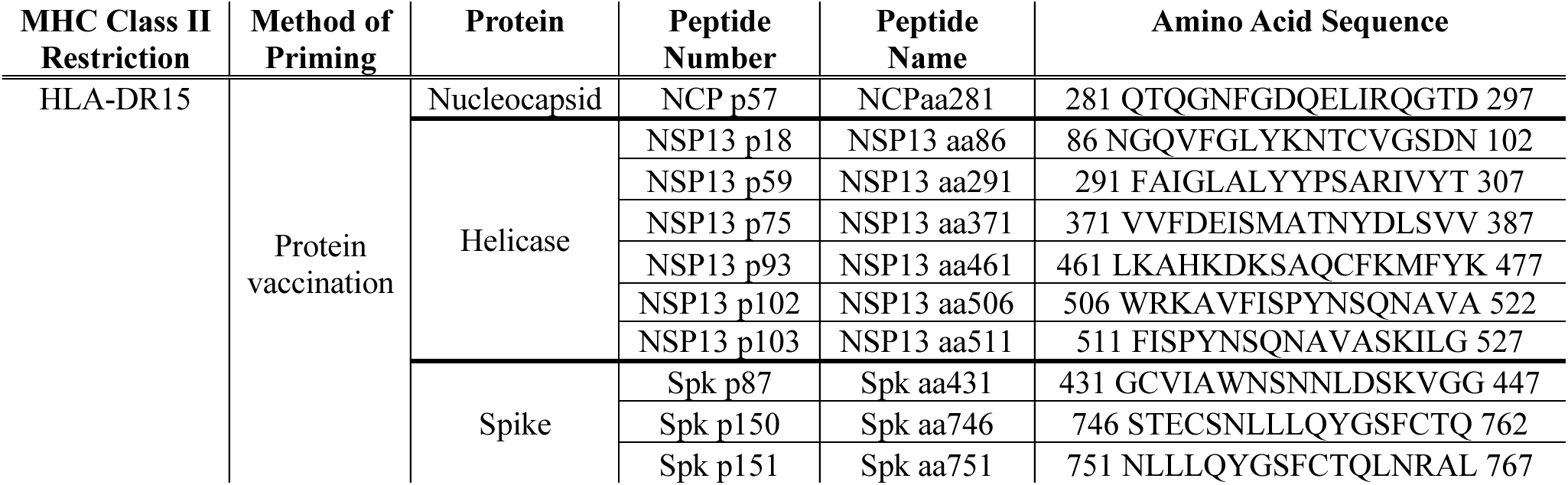
HLA-DR15 peptide sequences with amino acid numbers, peptide names and numbers used in the figures and the method of priming used to map the peptide epitopes in SARS-CoV-2.

In **Figure 8**, we illustrate the process and benefit of the systematic and unbiased approach we have described here for epitope discovery, which we call a funnel approach, an iterative process involving sequential elimination of many candidate epitopes based on direct *ex vivo* assays followed by determination of the immunogenic peptides. The top slice of the funnel (**A**) shows the potential number of peptide epitopes from all the proteins that were tested in each HLA-DR transgenic mouse strain, based on the number of overlapping peptides in each of the Influenza B (**Figure 8, top**) or SARS-CoV-2 (**Figure 8, bottom**) proteins. Some mice strains were not tested by IAV infection because of high pathogenicity. The sequential slices of the funnel indicate the number of peptides that remain at each stage after testing total pooled peptides, eliminating those in pools that elicited fewer than 50 spots (**B**). For large proteins (HA, NP and NA) that were positive based on total pools, peptide pooling matrices enabled further elimination of peptides based on non-stimulatory peptide pools from large proteins (**C**). These sequential processes left single candidate peptides from HA, NP and NA, as well as the complete set of single overlapping peptides from small proteins (NS1 and M1) that were tested individually, allowing further elimination of non-immunogenic peptides **(D)**. This process yielded the immunogenic, HLA-DR restricted peptide candidates that could then be used to test in HLA-typed human CD4 T cells. Depending on the HLA-DR allele, this process enriched candidates from 500-600 to approximately 5-24.

**Figure 8.**
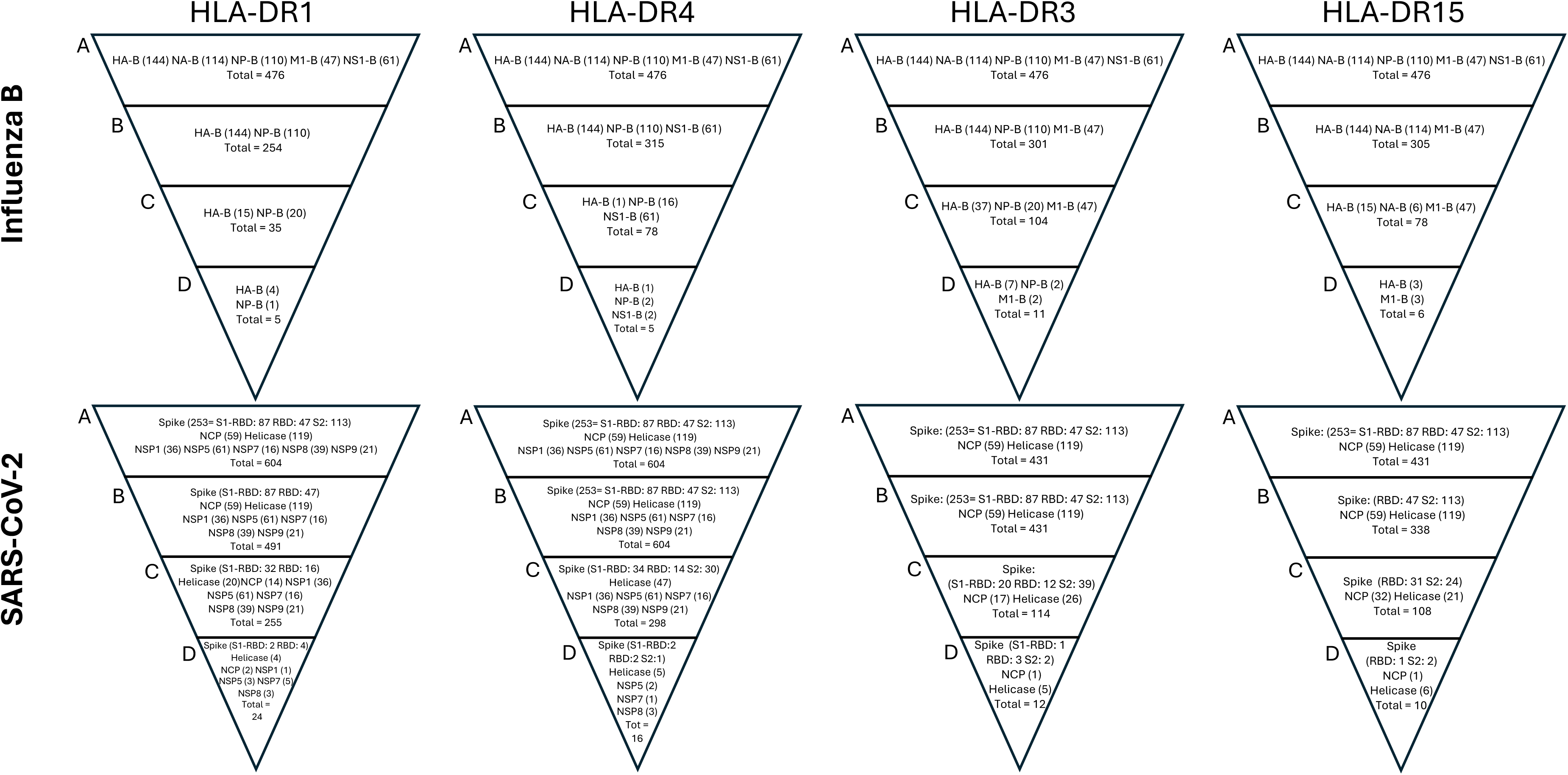
A funnel approach to CD4 T cell epitope identification. Shown are funnel diagrams of the iterative process of epitope discovery for influenza B (Top) or SARS-CoV-2 epitopes (bottom). In the top slice of each funnel (“A”) the total number of peptides screened for each individual HLA-DR allele or virus is indicated, with the proteins listed and the number of peptides contained in each protein shown in parathesis. In slice “B”, narrowing of the potential candidates using overlapping pools of peptides is shown. If no proteins were eliminated by pools either because pools were all positive or because pools were not used this value is the value in “A”. In slice “C” the potential single epitopes tested from either matrix or single peptides from small proteins such as NS1 or M1 is indicated. Finally in slice “D”, the number of positive peptides from the single peptide analysis is indicated. The individual peptides have been shown in the horizontal bar graphs in Figures 3 and 7 and listed in Tables I - VIII. These are the dominant CD4 peptide-epitope candidates for applications in humans or further animal models.

### Testing of reactivity in HLA-typed human subjects

After the comprehensive identification of peptide epitopes using the HLA-DR transgenic mice, we evaluated if these single peptides were sufficiently immunogenic in humans to be detected directly *ex vivo*, without any expansion in culture, a common strategy used by other groups ^52–54^, due to the low frequencies of single epitope-specific T cells from human PBMC. We surveyed the single HLA-DR restricted epitopes identified from circulating CD4 T cells from healthy adults with no known recent history of infection or vaccination, thus primarily reflecting the memory compartment. We focused on peptides that were dominant in the response to infection or vaccination in the HLA-Tg mice and that would be useful for human studies, such as those derived from surface receptors (influenza HA and SARS-CoV-2 spike), that are the target of vaccinations, or epitopes from internal virion proteins that are genetically conserved. It was anticipated that the immunodominance of any given epitope in humans, unlike the primary response studied in animal models, could be influenced by such factors as viral protein abundance after infection, persistence of epitope-specific CD4 T cells in memory or repeated boosting of T cells over time.

To assess single epitope-specific cells in humans, CD4 T cells were enriched from cryopreserved PBMC by depletion of CD8 and NK cells (which can also produce IFN-γ), leaving primarily APC and CD4 T cells. These CD4 T cell-enriched populations were tested with the single dominant peptides identified using HLA-DR transgenic mice. Approximately 9-12 HLA-typed donors were tested for each allele of HLA-DR. **Figure 9 (top)** shows the results with HLA-DR typed donors tested with influenza-derived peptide epitopes identified from N1, M1 and NP and HA proteins. Our data revealed that CD4 T cell epitope-specific cells varied among subjects, likely reflecting both the infection and vaccination history, as well as inherent subject-dependent magnitude and phenotype of responses shown to be predicted even before influenza vaccination ^55^. For example, some HLA-DR1 subjects exhibited relatively robust response to many epitopes, including those derived from influenza HA (HA-B aa271 p46/47) and M1 (M1 aa127 and aa205), and other subjects who were modest in the frequency of single peptide-specific CD4 T cells detected. When HLA-DR4 subjects, HLA-DR3 and HLA-DR15 were probed in a similar way, reactivity to single peptides derived from H1, H3, HA-B, M1, NP and NS1 were evaluated, many were also confirmed.

**Figure 9.**
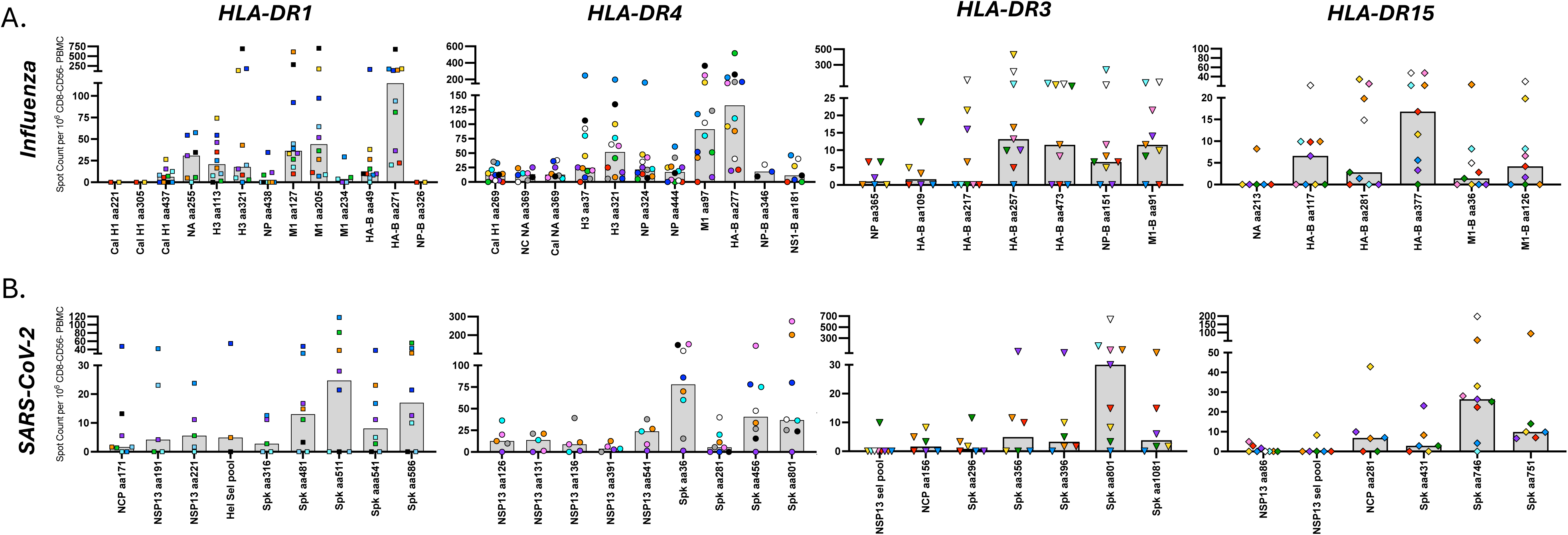
Human CD4 T responses to Influenza H1, H3 and B epitopes and SARS-CoV-2 epitopes. PBMCs from healthy adults that were collected between 2018 and 2023 and typed for HLA-DRB1*01:01 (far left), HLA-DRB1*04:01 (left middle), HLA-DRB1*03:01 (right middle) or HLA-DRB1*15:01 (far right) were enriched for CD4 T cells and APC by depletion of CD8 and CD56+ cells. CD4 T cell reactivity to each of the individual influenza derived peptides (top, A) and SARS-CoV-2 derived peptides (bottom, B) previously identified by epitope mapping in HLA-DR transgenic assessed in IFN-γ ELISpot assays is shown. Individual subjects are represented by unique symbols (Supplemental Table III), and the median response is shown as a grey bar. All data are represented as spots per million CD4 T cell-enriched PBMC, with background subtracted.

The ability of the SARS-CoV-2 derived peptides identified in HLA-DR transgenic mice to stimulate IFNγ production was then examined in human CD4 T cells **(Figure 9, bottom)**. Most donors exhibited detectable and sometimes robust reactivity to several peptides, particularly to spike-derived single peptides. For example, HLA-DR1 subjects have a robust response to spike aa511 and HLA-DR4 subjects to spike aa36. This is not surprising, as these specificities would be elicited by infection and vaccination. Interestingly, CD4 T cell reactivity was also detected to some non-spike epitopes, including N and Helicase, suggesting that these were induced by infection. Individuals displayed unique patterns with some peptides eliciting robust responses, while displaying little reactivity to others, indicating that the cytokine responses detected were epitope-specific. The HLA-DR restricted specificities identified here provided candidates for further studies.

### Tetramer staining enriches CD4 T cell memory populations from baseline blood samples

To further evaluate the funnel approach in identifying viral epitopes, we next determined if select peptide:HLA tetramers could enrich CD4 T cell responses directly from human PBMC. Tetramers were generated from three HLA-DR4-restricted peptides, including IAV H3 p81, IAV M1 p17, and SARS-CoV-2 S p8, were selected to evaluate CD4 responses across two common virus families. A set of negative control DR4 tetramers bound to the CLIP peptide were also generated. PBMCs from six healthy adult donors with confirmed HLA-DRB1*04:01 were then stained with tetramers and enriched using tetramer-associated magnetic enrichment (TAME). Tetramer staining with the DR4 CLIP tetramers indicated minimal background staining. Circulating CD4 T cells positive for IAV H3 p81 and M1 p17 were detected in all donor samples (**Figure 10, left**). SARS-CoV-2 S p8 tetramer-positive CD4 T cells were only detected in donors sampled after 2019 (Donors 38, 39, 40, 52), while no SARS-CoV-2 Sp8 tetramer-positive CD4 T cells were detected in pre-2019 Donors 17 and 31 (**Figure 10, left**). To determine the composition of the CD4 T cells enriched by tetramer staining, the tetramer-positive fractions and unenriched PBMC samples were surface stained with a CD4 memory phenotyping panel. Compared to the unenriched PBMCs, tetramer-positive CD4 T cells were enriched for central memory (TCM) and effector memory (TEM) populations while minimizing naive T cell staining in both frequency of TAME-enriched CD4 T cells (**Figure 10, left bar plot)** and number per million CD4 T cells (**Figure 10, right bar plot**). Together, these data support the funnel approach as a viable option for human virus epitope discovery that can enable generation of tetramers capable of enriching and analyzing CD4 T cell memory populations directly from donor PBMCs.

**Figure 10.**
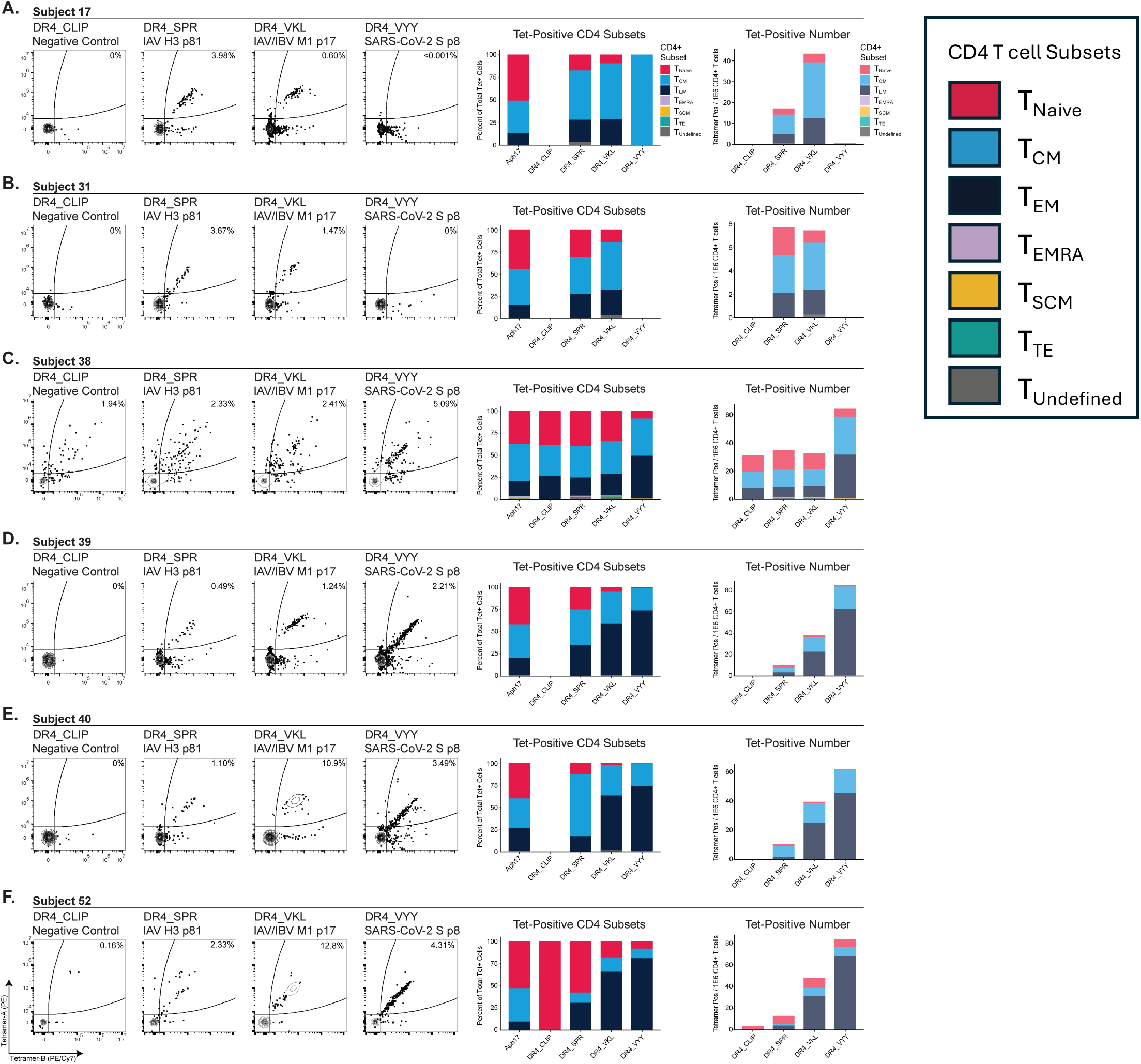
Characterization of human CD4+ T subsets following tetramer-associated magnetic enrichment. (A-F) PBMCs from healthy adult donors with HLA-DRB1*04:01 were dual-stained with PE- and *PE/Cy7-conjugated HLA-DR4 tetramers loaded with CLIP (negative control*), IAV H3 p81 (SPRYVKQNTLKLATGMR), IAV/IBV M1 p17 (VKLYRKLKREITFHGAK), or SARS-CoV-2 S p8 (VYYPDKVFRSSVLHSTQ) and subjected to tetramer-associated magnetic enrichment (TAME). TAME-selected cells were surface stained with CD4 immunophenotyping panel and analyzed by flow cytometry. Flow plots depict percentages of cells positive for tetramer on both colors (Tet+) as a percentage of TAME-selected (top; inset) and total (bottom; inset) live CD3+CD4+ lymphocytes. The composition of Tet+ CD4+ T cell subsets are depicted in the bar plots as either the percentage of total Tet+ cells for a given tetramer (middle) or Tet+ counts per 1×10^6^ CD4+ T cells (far right). The composition of the total CD4+ populations (non-TAME) for each subject are provided (left bar). Bar plot positions with no values indicate that no Tet+ cells were identified. CD4+ T cell subsets included T_Naïve_ (naïve); T_SCM_ (stem cell memory); T_CM_ (central memory); T_EM_ (effector memory); T_EMRA_ (effector memory CD45RA positive); T_TE_ (terminally exhausted); T_Undefined_ (not defined by other categories listed). Flow plots and corresponding bar plots represent individually stained aliquots of PBMCs from each subject.

## Discussion

In this study, we defined a diverse set of CD4 T cell epitopes from the respiratory pathogens Influenza A, Influenza B, and SARS-CoV-2. The goal of these studies was to define candidate epitopes to further research in the field of human CD4 T cell immunity and immune memory generated by pathogen infection and vaccination. Often CD4 T cell immunity is evaluated within our own laboratory ^51,56–58^ as well as others ^59–62^, using large peptide pools. Although very useful to “capture” the full repertoire of antigen-specific CD4 T cells from known pathogen derived proteins, these approaches are limited for end stage analyses because they cannot resolve the heterogeneity in reactivity to particular types of epitopes or enable approaches such as T cell receptor repertoire analyses or gene expression profiling. Single epitope identification enables multimer-based isolation of CD4 T cells for transcriptional profiling and for tracking the trajectory and diversity of the T cell repertoire over time in humans ^63–65^.

We did note some differences in the immunodominance of peptides identified in the animal studies and those determined from HLA-typed human donors. Reactivity in humans will likely reflect not only inherent differences in immunogenicity but variability in the number of confrontations humans have experienced over their lifetime and thus may become enriched for more conserved epitopes. Also, competition in CD4 T cell responses among alternate HLA-class II molecules in the typical human host may modulate the relative immunodominance of the particular peptide:HLA-DR tested ^66,67^. Finally, immune responses in HLA-DR transgenic mice may be modulated by suboptimal interactions between the human HLA-DR protein and the mouse peptide editor DM (H-2M). CD4 T cell interactions with class II as a “co-receptor” may be suboptimal in DR-transgenic mice, as only some (HLA-DR1 and HLA-DR4) incorporated the mouse β2 domain into the HLA-DR transgene, the region of MHC-class II which has been identified as a dominant site for CD4 T cell interaction ^68^. Thus, the quantitation presented here is best used as a positive empirically defined indicator of the immunogenicity of epitope-specific CD4 T cells that have the potential to be elicited in humans after infection or vaccination. Subjects of different ages, varying histories of infection and/or vaccination and different arrays of co-expressed HLA-class II alleles will likely display variability in CD4 T cell epitope prominence.

There are several limitations in this study. First, some proteins in influenza have not been screened for epitopes with HLA-DR transgenic mice, because of pathogenicity of some of the viruses in the HLA-DR transgenic mice. For example, the polymerase protein epitopes were not tested, although we have found that this protein can be immunogenic in several inbred strains of mice ^69,70^ and could thus contain highly conserved CD4 T cell epitopes. Second, we were limited in the targets from SARS-CoV-2 that could be mapped for CD4 T cell epitopes due to limited availability recombinant proteins for vaccination. Third, the relative reactivity in the human samples was drawn from limited subjects, so the positive response is informative but the magnitude less so. Finally, we did not selectively recruit HLA-typed subjects who have a history of recent infection, and thus the relative immunodominance across epitopes might shift in a given subject pre and post infection. However, our ability to detect and characterize the epitope specific CD4 T cells in resting memory cells, without *in vitro* expansion, does demonstrate the fidelity of the approach we have adopted here.

## Acknowledgements

We would like to thank members of A. Joachimiak lab, especially Dr. Robert Jedrzejczak who cloned and purified SARS-CoV-2 proteins. This work was supported with Federal funds from the National Institute of Allergy and Infectious Diseases, National Institutes of Health, Department of Health and Human Services, U01 AI14461602S1 and Centers of Excellence for Influenza Research and Response (CEIRR-CIDER) contract number 75N93021C00018 sub-contracts to AJS. This work was supported by Federal funds from the National Institute of Allergy and Infectious Diseases, National Institutes of Health, Department of Health and Human Services, under Contracts HHSN272201700060C and 75N93022C00035 (AJ), by the DOE Office of Science through the National Virtual Biotechnology Laboratory, a consortium of DOE national laboratories focused on response to COVID-19, with funding provided by the Coronavirus CARES Act (AJ). This work was supported with Federal funds from the National Institute of Allergy and Infectious Diseases, National Institutes of Health, Department of Health and Human Services, U01 AI14461602S1 to PT. RCM is supported by a Ruth L. Kirschstein National Research Service Award (NRSA) Individual Postdoctoral Fellowship (NRSA-NIAID) F32AI157296. The content of this article is solely the responsibility of the authors and does not necessarily represent the official views of the National Institutes of Health.

## Data Availability

The full complement of data accumulated for these studies is available upon request.

---

**Supplemental Table I.** Detailed information for peptide arrays.

| Virus | Protein | Number of Peptides | Length of peptide | Amino Acid Overlap | Source | BEI Resources (BEIR) Reference Number |
| --- | --- | --- | --- | --- | --- | --- |
| H1N1<br>A/California/04/2009 | HA | 139 | 15 | 11 | BEIR | NR-15433 |
|  | NA | 115 | 13-16 | 10-12 | BEIR | NR-18975 |
|  | NP | 122 | 14-15 | 11 | BEIR | NR-18976 |
|  | M1 | 61 | 11-16 | 10-12 | BEIR | NR-18977 |
|  | NS1 | 52 | 14-17 | 10-13 | BEIR | NR-18979 |
| H1N1<br>A/New<br>Caledonia/20/1999 | HA | 94 | 17 | 11 | BEIR | NR-2602 |
|  | NA | 78 | 17 | 11 | BEIR | NR-2606 |
| H1N1<br>A/New York/348/2003 | NP | 82 | 17 | 11 | BEIR | NR-2611 |
|  | M1 | 41 | 17 | 11 | BEIR | NR-2613 |
| H3N2 A/Perth/16/2009 | HA | 139 | 15 | 11 | BEIR | NR-19266 |
| IBV<br>B/Brisbane/60/2008 | HA | 144 | 13-15 | 11 | BEIR | NR-19247 |
|  | NA | 114 | 14-15 | 11 | BEIR | NR-19523 |
| IBV<br>B/Florida/04/2006 | HA | 96 | 17 | 11 | BEIR | NR-18972 |
|  | NA | 114 | 13-16 | 10-12 | BEIR | NR-19254 |
|  | NP | 110 | 15-17 | 11 | BEIR | NR-36045 |
|  | M1 | 47 | 17 | 10-12 | BEIR | NR-36046 |
|  | NS1 | 68 | 17 | 11 | Mimotopes |  |
| SARS-CoV-2 | NCP | 59 | 17 or 13 | 10 | BEIR | NR-52404 |
|  | Spike | 253 | 17 | 11 | Mimotopes |  |
|  | Helicase | 119 | 17 | 11 | Mimotopes |  |
|  | NSP1 | 36 | 17 | 11 | Mimotopes |  |
|  | NSP5 | 61 | 17 | 11 | Mimotopes |  |
|  | NSP7 | 16 | 17 | 11 | Mimotopes |  |
|  | NSP8 | 39 | 17 | 11 | Mimotopes |  |
|  | NSP9 | 21 | 17 | 11 | Mimotopes |  |
|  | NSP15 | 67 | 17 | 11 | Mimotopes |  |

**Supplemental Table II.** Antibodies used in tetramer staining panel.

| Antibody | Fluorochrome | Clone | Dilution | Supplier | Item Identifier |
| --- | --- | --- | --- | --- | --- |
| CD95 | BUV395 | DX2 | 1:20 | BD Biosciences | 740306 |
| CD62L | BUV496 | SK11 | 1:20 | BD Biosciences | 750589 |
| CD28 | BUV563 | CD28.2 | 1:20 | BD Biosciences | 741392 |
| CD45RA | BV421 | HI100 | 1:20 | Biolegend | 304130 |
| CD25 | V450 | M-A251 | 1:20 | BD Biosciences | 560355 |
| CD45RO | BV605 | UCHL1 | 1:20 | Biolegend | 304238 |
| CD4 | BV650 | SK3 | 1:20 | Biolegend | 344692 |
| CD11a | BV711 | HI111 | 1:20 | Biolegend | 301238 |
| CD27 | BV785 | O323 | 1:20 | Biolegend | 302832 |
| CCR7 | FITC | G043H7 | 1:20 | Biolegend | 353216 |
| CD58 | PerCP-Cy5.5 | TS2/9 | 1:20 | Biolegend | 330914 |
| CD3 | PerCP-eFluor 710 | OKT3 | 1:20 | ThermoFisher | 46-0037-42 |
| NKG2D | APC | 1D11 | 1:20 | Biolegend | 320808 |
| CXCR3 | AF700 | G025H7 | 1:20 | Biolegend | 353742 |
| Live/Dead | BV510 |  | 1:400 | Tonbo | 13-0870-T500 |

**Supplemental Table III.**
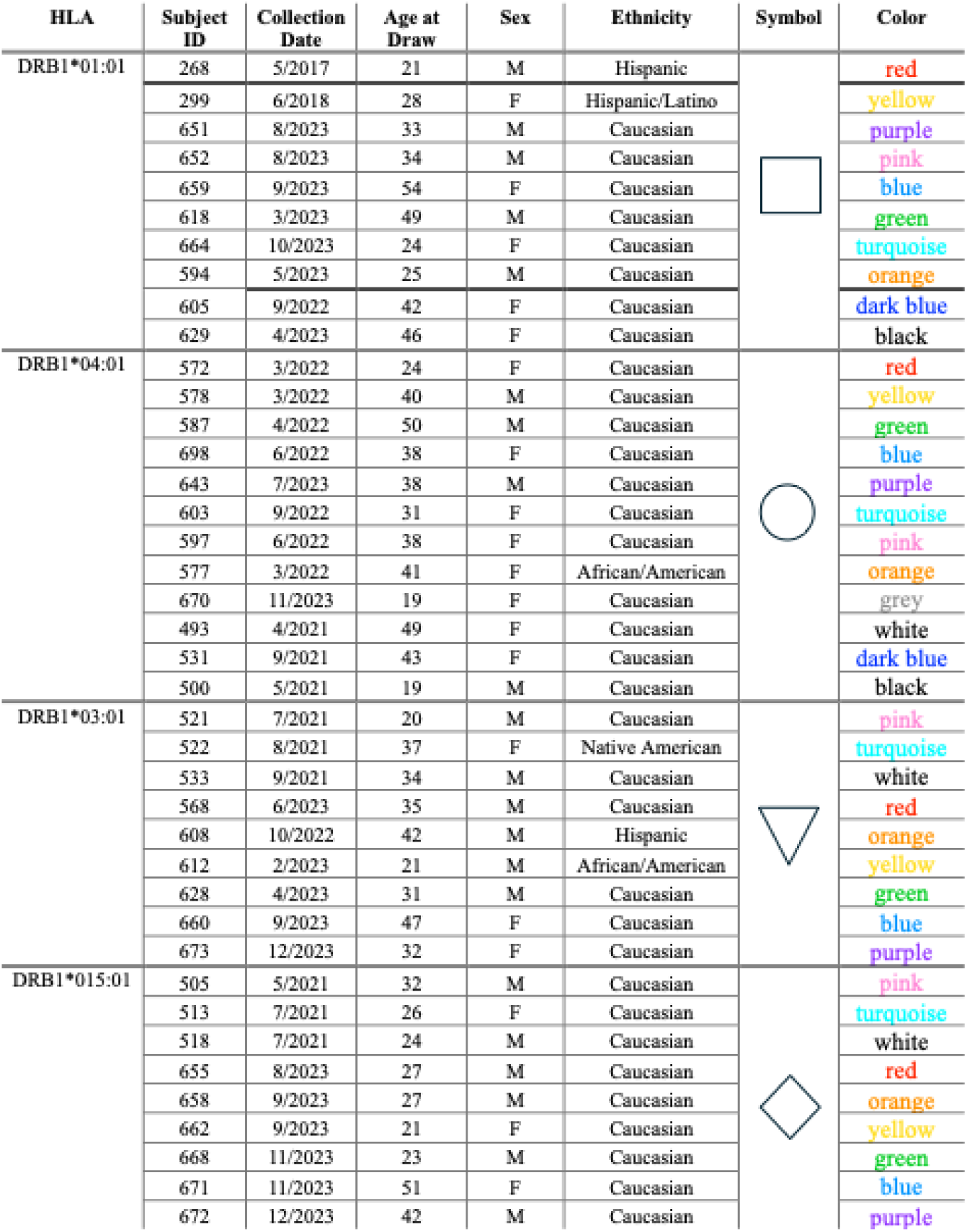
Demographics information and unique symbol identifiers for PBMC samples from healthy adult subjects that were purchased from CTL and used in Figure 9.

## References

1. Altman, J.D., and Davis, M.M. (2016). MHC-Peptide Tetramers to Visualize Antigen-Specific T Cells. Curr Protoc Immunol 115, 17 13 11–17 13 44. 10.1002/cpim.14.

2. Tiwari, R., Singh, V.K., Rajneesh, Kumar, A., Gautam, V., and Kumar, R. (2024). MHC tetramer technology: Exploring T cell biology in health and disease. Adv Protein Chem Struct Biol 140, 327–345. 10.1016/bs.apcsb.2024.02.002.

3. Massilamany, C., Krishnan, B., and Reddy, J. (2015). Major Histocompatibility Complex Class II Dextramers: New Tools for the Detection of antigen-Specific, CD4 T Cells in Basic and Clinical Research. Scand J Immunol 82, 399–408. 10.1111/sji.12344.

4. Reber, A.J., Music, N., Kim, J.H., Gansebom, S., Chen, J., and York, I. (2018). Extensive T cell cross-reactivity between diverse seasonal influenza strains in the ferret model. Sci Rep 8, 6112. 10.1038/s41598-018-24394-z.

5. Roti, M., Yang, J., Berger, D., Huston, L., James, E.A., and Kwok, W.W. (2008). Healthy human subjects have CD4+ T cells directed against H5N1 influenza virus. J Immunol 180, 1758–1768.

6. Richards, K.A., Chaves, F.A., Krafcik, F.R., Topham, D.J., Lazarski, C.A., and Sant, A.J. (2007). Direct ex vivo analyses of HLA-DR1 transgenic mice reveal an exceptionally broad pattern of immunodominance in the primary HLA-DR1-restricted CD4 T-cell response to influenza virus hemagglutinin. J Virol 81, 7608–7619. 10.1128/JVI.02834-06.

7. Greenbaum, J.A., Kotturi, M.F., Kim, Y., Oseroff, C., Vaughan, K., Salimi, N., Vita, R., Ponomarenko, J., Scheuermann, R.H., Sette, A., and Peters, B. (2009). Pre-existing immunity against swine-origin H1N1 influenza viruses in the general human population. Proc Natl Acad Sci U S A 106, 20365–20370. 10.1073/pnas.0911580106.

8. Redd, A.D., Nardin, A., Kared, H., Bloch, E.M., Abel, B., Pekosz, A., Laeyendecker, O., Fehlings, M., Quinn, T.C., and Tobian, A.A.R. (2022). Minimal Crossover between Mutations Associated with Omicron Variant of SARS-CoV-2 and CD8(+) T-Cell Epitopes Identified in COVID-19 Convalescent Individuals. mBio 13, e0361721. 10.1128/mbio.03617-21.

9. Tarke, A., Coelho, C.H., Zhang, Z., Dan, J.M., Yu, E.D., Methot, N., Bloom, N.I., Goodwin, B., Phillips, E., Mallal, S., et al. (2022). SARS-CoV-2 vaccination induces immunological T cell memory able to cross-recognize variants from Alpha to Omicron. Cell 185, 847–859 e811. 10.1016/j.cell.2022.01.015.

10. Tan, P.T., Heiny, A.T., Miotto, O., Salmon, J., Marques, E.T., Lemonnier, F., and August, J.T. (2010). Conservation and diversity of influenza A H1N1 HLA-restricted T cell epitope candidates for epitope-based vaccines. PLoS One 5, e8754. 10.1371/journal.pone.0008754.

11. Schroeder, S.M., Nelde, A., and Walz, J.S. (2023). Viral T-cell epitopes - Identification, characterization and clinical application. Semin Immunol 66, 101725. 10.1016/j.smim.2023.101725.

12. Muraduzzaman, A.K.M., Illing, P.T., Mifsud, N.A., and Purcell, A.W. (2022). Understanding the Role of HLA Class I Molecules in the Immune Response to Influenza Infection and Rational Design of a Peptide-Based Vaccine. Viruses 14. 10.3390/v14112578.

13. Lenz, T.L. (2024). HLA Genes: A Hallmark of Functional Genetic Variation and Complex Evolution. Methods Mol Biol 2809, 1–18. 10.1007/978-1-0716-3874-3_1.

14. Nayak, J.L., Richards, K.A., Chaves, F.A., and Sant, A.J. (2010). Analyses of the specificity of CD4 T cells during the primary immune response to influenza virus reveals dramatic MHC-linked asymmetries in reactivity to individual viral proteins. Viral Immunol 23, 169–180. 10.1089/vim.2009.0099.

15. DiPiazza, A., Richards, K., Poulton, N., and Sant, A.J. (2017). Avian and Human Seasonal Influenza Hemagglutinin Proteins Elicit CD4 T Cell Responses That Are Comparable in Epitope Abundance and Diversity. Clin Vaccine Immunol 24. 10.1128/CVI.00548-16.

16. Tobery, T.W., and Caulfield, M.J. (2004). Identification of T-cell epitopes using ELISpot and peptide pool arrays. Methods Mol Med 94, 121–132. 10.1385/1-59259-679-7:121.

17. Peters, B., Nielsen, M., and Sette, A. (2020). T Cell Epitope Predictions. Annu Rev Immunol 38, 123–145. 10.1146/annurev-immunol-082119-124838.

18. Archila, L.L., and Kwok, W.W. (2017). Tetramer-Guided Epitope Mapping: A Rapid Approach to Identify HLA-Restricted T-Cell Epitopes from Composite Allergens. Methods Mol Biol 1592, 199–209. 10.1007/978-1-4939-6925-8_16.

19. Sidney, J., Peters, B., and Sette, A. (2020). Epitope prediction and identification-adaptive T cell responses in humans. Semin Immunol 50, 101418. 10.1016/j.smim.2020.101418.

20. Clement, C.C., Becerra, A., Yin, L., Zolla, V., Huang, L., Merlin, S., Follenzi, A., Shaffer, S.A., Stern, L.J., and Santambrogio, L. (2016). The Dendritic Cell Major Histocompatibility Complex II (MHC II) Peptidome Derives from a Variety of Processing Pathways and Includes Peptides with a Broad Spectrum of HLA-DM Sensitivity. J Biol Chem 291, 5576–5595. 10.1074/jbc.M115.655738.

21. Yang, Y., Wei, Z., Cia, G., Song, X., Pucci, F., Rooman, M., Xue, F., and Hou, Q. (2024). MHCII- peptide presentation: an assessment of the state-of-the-art prediction methods. Front Immunol 15, 1293706. 10.3389/fimmu.2024.1293706.

22. Feola, S., Chiaro, J., and Cerullo, V. (2023). Integrating immunopeptidome analysis for the design and development of cancer vaccines. Semin Immunol 67, 101750. 10.1016/j.smim.2023.101750.

23. Ahn, R., Cui, Y., and White, F.M. (2023). Antigen discovery for the development of cancer immunotherapy. Semin Immunol 66, 101733. 10.1016/j.smim.2023.101733.

24. Illing, P.T., Ramarathinam, S.H., and Purcell, A.W. (2022). New insights and approaches for analyses of immunopeptidomes. Curr Opin Immunol 77, 102216. 10.1016/j.coi.2022.102216.

25. Becker, J.P., and Riemer, A.B. (2022). The Importance of Being Presented: Target Validation by Immunopeptidomics for Epitope-Specific Immunotherapies. Front Immunol 13, 883989. 10.3389/fimmu.2022.883989.

26. Chong, C., Coukos, G., and Bassani-Sternberg, M. (2022). Identification of tumor antigens with immunopeptidomics. Nat Biotechnol 40, 175–188. 10.1038/s41587-021-01038-8.

27. Woon, A.P., and Purcell, A.W. (2018). The use of proteomics to understand antiviral immunity. Semin Cell Dev Biol 84, 22–29. 10.1016/j.semcdb.2017.12.002.

28. Joyce, S., and Ternette, N. (2021). Know thy immune self and non-self: Proteomics informs on the expanse of self and non-self, and how and where they arise. Proteomics 21, e2000143. 10.1002/pmic.202000143.

29. Lazarski, C.A., Chaves, F.A., Jenks, S.A., Wu, S., Richards, K.A., Weaver, J.M., and Sant, A.J. (2005). The kinetic stability of MHC class II:peptide complexes is a key parameter that dictates immunodominance. Immunity 23, 29–40. 10.1016/j.immuni.2005.05.009.

30. Rasmussen, M., Fenoy, E., Harndahl, M., Kristensen, A.B., Nielsen, I.K., Nielsen, M., and Buus, S. (2016). Pan-Specific Prediction of Peptide-MHC Class I Complex Stability, a Correlate of T Cell Immunogenicity. J Immunol 197, 1517–1524. 10.4049/jimmunol.1600582.

31. Hensen, L., Illing, P.T., Rowntree, L.C., Davies, J., Miller, A., Tong, S.Y.C., Habel, J.R., van de Sandt, C.E., Flanagan, K.L., Purcell, A.W., et al. (2022). T Cell Epitope Discovery in the Context of Distinct and Unique Indigenous HLA Profiles. Front Immunol 13, 812393. 10.3389/fimmu.2022.812393.

32. Karlsson, L. (2005). DM and DO shape the repertoire of peptide-MHC-class-II complexes. Curr Opin Immunol 17, 65–70. 10.1016/j.coi.2004.11.003.

33. Sant, A.J., Chaves, F.A., Jenks, S.A., Richards, K.A., Menges, P., Weaver, J.M., and Lazarski, C.A. (2005). The relationship between immunodominance, DM editing, and the kinetic stability of MHC class II:peptide complexes. Immunol Rev 207, 261–278. 10.1111/j.0105-2896.2005.00307.x.

34. van Hateren, A., and Elliott, T. (2023). Visualising tapasin- and TAPBPR-assisted editing of major histocompatibility complex class-I immunopeptidomes. Curr Opin Immunol 83, 102340. 10.1016/j.coi.2023.102340.

35. Thomas, C., and Tampe, R. (2021). MHC I assembly and peptide editing - chaperones, clients, and molecular plasticity in immunity. Curr Opin Immunol 70, 48–56. 10.1016/j.coi.2021.02.004.

36. Welsh, R.A., and Sadegh-Nasseri, S. (2020). The love and hate relationship of HLA-DM/DO in the selection of immunodominant epitopes. Curr Opin Immunol 64, 117–123. 10.1016/j.coi.2020.05.007.

37. Jenkins, M.K., and Moon, J.J. (2012). The role of naive T cell precursor frequency and recruitment in dictating immune response magnitude. J Immunol 188, 4135–4140. 10.4049/jimmunol.1102661.

38. Tscharke, D.C., Croft, N.P., Doherty, P.C., and La Gruta, N.L. (2015). Sizing up the key determinants of the CD8(+) T cell response. Nat Rev Immunol 15, 705–716. 10.1038/nri3905.

39. Turner, S.J., Kedzierska, K., Komodromou, H., La Gruta, N.L., Dunstone, M.A., Webb, A.I., Webby, R., Walden, H., Xie, W., McCluskey, J., et al. (2005). Lack of prominent peptide-major histocompatibility complex features limits repertoire diversity in virus-specific CD8+ T cell populations. Nat Immunol 6, 382–389. 10.1038/ni1175.

40. Rosloniec, E.F., Brand, D.D., Myers, L.K., Whittington, K.B., Gumanovskaya, M., Zaller, D.M., Woods, A., Altmann, D.M., Stuart, J.M., and Kang, A.H. (1997). An HLA-DR1 transgene confers susceptibility to collagen-induced arthritis elicited with human type II collagen. J Exp Med 185, 1113–1122. 10.1084/jem.185.6.1113.

41. Woods, A., Chen, H.Y., Trumbauer, M.E., Sirotina, A., Cummings, R., and Zaller, D.M. (1994). Human major histocompatibility complex class II-restricted T cell responses in transgenic mice. J Exp Med 180, 173–181. 10.1084/jem.180.1.173.

42. Vandenbark, A.A., Rich, C., Mooney, J., Zamora, A., Wang, C., Huan, J., Fugger, L., Offner, H., Jones, R., and Burrows, G.G. (2003). Recombinant TCR ligand induces tolerance to myelin oligodendrocyte glycoprotein 35-55 peptide and reverses clinical and histological signs of chronic experimental autoimmune encephalomyelitis in HLA-DR2 transgenic mice. J Immunol 171, 127–133. 10.4049/jimmunol.171.1.127.

43. Strauss, G., Vignali, D.A., Schonrich, G., and Hammerling, G.J. (1994). Negative and positive selection by HLA-DR3(DRw17) molecules in transgenic mice. Immunogenetics 40, 104–108.

44. Cosgrove, D., Gray, D., Dierich, A., Kaufman, J., Lemeur, M., Benoist, C., and Mathis, D. (1991). Mice lacking MHC class II molecules. Cell 66, 1051–1066. 10.1016/0092-8674(91)90448-8.

45. Tobery, T.W., Wang, S., Wang, X.M., Neeper, M.P., Jansen, K.U., McClements, W.L., and Caulfield, M.J. (2001). A simple and efficient method for the monitoring of antigen-specific T cell responses using peptide pool arrays in a modified ELISpot assay. J Immunol Methods 254, 59–66. 10.1016/s0022-1759(01)00397-0.

46. Pascua, P.N.Q., Mostafa, H.H., Marathe, B.M., Vogel, P., Russell, C.J., Webby, R.J., and Govorkova, E.A. (2017). Pathogenicity and peramivir efficacy in immunocompromised murine models of influenza B virus infection. Sci Rep 7, 7345. 10.1038/s41598-017-07433-z.

47. Kirkpatrick Roubidoux, E., McMahon, M., Carreno, J.M., Capuano, C., Jiang, K., Simon, V., van Bakel, H., Wilson, P., and Krammer, F. (2021). Identification and Characterization of Novel Antibody Epitopes on the N2 Neuraminidase. mSphere 6. 10.1128/mSphere.00958-20.

48. Rattan, A., White, C.L., Nelson, S., Eismann, M., Padilla-Quirarte, H., Glover, M.A., Dileepan, T., Marathe, B.M., Govorkova, E.A., Webby, R.J., et al. (2022). Development of a Mouse Model to Explore CD4 T Cell Specificity, Phenotype, and Recruitment to the Lung after Influenza B Infection. Pathogens 11. 10.3390/pathogens11020251.

49. Nguyen-Contant, P., Embong, A.K., Kanagaiah, P., Chaves, F.A., Yang, H., Branche, A.R., Topham, D.J., and Sangster, M.Y. (2020). S Protein-Reactive IgG and Memory B Cell Production after Human SARS-CoV-2 Infection Includes Broad Reactivity to the S2 Subunit. mBio 11. 10.1128/mBio.01991-20.

50. Guthmiller, J.J., Stovicek, O., Wang, J., Changrob, S., Li, L., Halfmann, P., Zheng, N.Y., Utset, H., Stamper, C.T., Dugan, H.L., et al. (2021). SARS-CoV-2 Infection Severity Is Linked to Superior Humoral Immunity against the Spike. mBio 12. 10.1128/mBio.02940-20.

51. Richards, K.A., Glover, M., Crawford, J.C., Thomas, P.G., White, C., and Sant, A.J. (2021). Circulating CD4 T Cells Elicited by Endemic Coronaviruses Display Vast Disparities in Abundance and Functional Potential Linked to Antigen Specificity and Age. J Infect Dis 223, 1555–1563. 10.1093/infdis/jiab076.

52. Reijonen, H., and Kwok, W.W. (2003). Use of HLA class II tetramers in tracking antigen-specific T cells and mapping T-cell epitopes. Methods 29, 282–288. 10.1016/s1046-2023(02)00350-x.

53. Michelo, C.M., Dalel, J.A., Hayes, P., Fernandez, N., Fiore-Gartland, A., Kilembe, W., Tang, J., Streatfield, C., Gilmour, J., and Hunter, E. (2021). Comprehensive epitope mapping using polyclonally expanded human CD8 T cells and a two-step ELISpot assay for testing large peptide libraries. J Immunol Methods 491, 112970. 10.1016/j.jim.2021.112970.

54. Mateus, J., Grifoni, A., Tarke, A., Sidney, J., Ramirez, S.I., Dan, J.M., Burger, Z.C., Rawlings, S.A., Smith, D.M., Phillips, E., et al. (2020). Selective and cross-reactive SARS-CoV-2 T cell epitopes in unexposed humans. Science 370, 89–94. 10.1126/science.abd3871.

55. Tsang, J.S., Schwartzberg, P.L., Kotliarov, Y., Biancotto, A., Xie, Z., Germain, R.N., Wang, E., Olnes, M.J., Narayanan, M., Golding, H., et al. (2014). Global analyses of human immune variation reveal baseline predictors of postvaccination responses. Cell 157, 499–513. 10.1016/j.cell.2014.03.031.

56. Richards, K.A., Changrob, S., Thomas, P.G., Wilson, P.C., and Sant, A.J. (2024). Lack of memory recall in human CD4 T cells elicited by the first encounter with SARS-CoV-2. iScience 27, 109992. 10.1016/j.isci.2024.109992.

57. Moritzky, S.A., Richards, K.A., Glover, M.A., Krammer, F., Chaves, F.A., Topham, D.J., Branche, A., Nayak, J.L., and Sant, A.J. (2023). The Negative Effect of Preexisting Immunity on Influenza Vaccine Responses Transcends the Impact of Vaccine Formulation Type and Vaccination History. J Infect Dis 227, 381–390. 10.1093/infdis/jiac068.

58. Richards, K.A., Treanor, J.J., Nayak, J.L., and Sant, A.J. (2018). Overarching Immunodominance Patterns and Substantial Diversity in Specificity and Functionality in the Circulating Human Influenza A and B CD4 T Cell Repertoire. J Infect Dis. 10.1093/infdis/jiy288.

59. Draenert, R., Altfeld, M., Brander, C., Basgoz, N., Corcoran, C., Wurcel, A.G., Stone, D.R., Kalams, S.A., Trocha, A., Addo, M.M., et al. (2003). Comparison of overlapping peptide sets for detection of antiviral CD8 and CD4 T cell responses. J Immunol Methods 275, 19–29. 10.1016/s0022-1759(02)00541-0.

60. Becerra-Artiles, A., Calvo-Calle, J.M., Co, M.D., Nanaware, P.P., Cruz, J., Weaver, G.C., Lu, L., Forconi, C., Finberg, R.W., Moormann, A.M., and Stern, L.J. (2022). Broadly recognized, cross-reactive SARS-CoV-2 CD4 T cell epitopes are highly conserved across human coronaviruses and presented by common HLA alleles. Cell Rep 39, 110952. 10.1016/j.celrep.2022.110952.

61. da Silva Antunes, R., Weiskopf, D., Sidney, J., Rubiro, P., Peters, B., Lindestam Arlehamn, C.S., Grifoni, A., and Sette, A. (2023). The MegaPool Approach to Characterize Adaptive CD4+ and CD8+ T Cell Responses. Curr Protoc 3, e934. 10.1002/cpz1.934.

62. Zhang, Z., Mateus, J., Coelho, C.H., Dan, J.M., Moderbacher, C.R., Galvez, R.I., Cortes, F.H., Grifoni, A., Tarke, A., Chang, J., et al. (2022). Humoral and cellular immune memory to four COVID-19 vaccines. Cell 185, 2434–2451 e2417. 10.1016/j.cell.2022.05.022.

63. Davis, M.M., and Boyd, S.D. (2019). Recent progress in the analysis of alphabetaT cell and B cell receptor repertoires. Curr Opin Immunol 59, 109–114. 10.1016/j.coi.2019.05.012.

64. Rowntree, L.C., Nguyen, T.H.O., Kedzierski, L., Neeland, M.R., Petersen, J., Crawford, J.C., Allen, L.F., Clemens, E.B., Chua, B., McQuilten, H.A., et al. (2022). SARS-CoV-2-specific T cell memory with common TCRalphabeta motifs is established in unvaccinated children who seroconvert after infection. Immunity 55, 1299–1315 e1294. 10.1016/j.immuni.2022.06.003.

65. Pachnio, A., Ciaurriz, M., Begum, J., Lal, N., Zuo, J., Beggs, A., and Moss, P. (2016). Cytomegalovirus Infection Leads to Development of High Frequencies of Cytotoxic Virus-Specific CD4+ T Cells Targeted to Vascular Endothelium. PLoS Pathog 12, e1005832. 10.1371/journal.ppat.1005832.

66. Smith, A.L., Wikstrom, M.E., and Fazekas de St Groth, B. (2000). Visualizing T cell competition for peptide/MHC complexes: a specific mechanism to minimize the effect of precursor frequency. Immunity 13, 783–794. 10.1016/s1074-7613(00)00076-5.

67. Hayball, J.D., Robinson, B.W., and Lake, R.A. (2004). CD4+ T cells cross-compete for MHC class II-restricted peptide antigen complexes on the surface of antigen presenting cells. Immunol Cell Biol 82, 103–111. 10.1046/j.0818-9641.2004.01233.x.

68. Konig, R., Huang, L.Y., and Germain, R.N. (1992). MHC class II interaction with CD4 mediated by a region analogous to the MHC class I binding site for CD8. Nature 356, 796–798. 10.1038/356796a0.

69. Alam, S., Chan, C., Qiu, X., Shannon, I., White, C.L., Sant, A.J., and Nayak, J.L. (2017). Selective pre-priming of HA-specific CD4 T cells restores immunological reactivity to HA on heterosubtypic influenza infection. PLoS One 12, e0176407. 10.1371/journal.pone.0176407.

70. Dunn-Walters, D., Townsend, C., Sinclair, E., and Stewart, A. (2018). Immunoglobulin gene analysis as a tool for investigating human immune responses. Immunol Rev 284, 132–147. 10.1111/imr.12659.

